# Probing the neural dynamics of mnemonic representations after the initial consolidation

**DOI:** 10.1101/803718

**Authors:** Wei Liu, Nils Kohn, Guillén Fernández

## Abstract

Memories are not stored as static engrams, but as dynamic representations affected by processes occurring after initial encoding. Previous studies revealed changes in activity and mnemonic representations in visual processing areas, parietal lobe, and hippocampus underlying repeated retrieval and suppression. However, these neural changes are usually induced by memory modulation immediately after memory formation. Here, we investigated 27 healthy participants with a two-day functional Magnetic Resonance Imaging design to probe how established memories are dynamically modulated by retrieval and suppression 24 hours after learning. Behaviorally, we demonstrated that established memories can still be strengthened by repeated retrieval. By contrast, repeated suppression had a modest negative effect, and suppression-induced forgetting was associated with individual suppression efficacy. Neurally, we demonstrated item-specific pattern reinstatements in visual processing areas, parietal lobe, and hippocampus. Then, we showed that repeated retrieval reduced activity amplitude in the ventral visual cortex and hippocampus, but enhanced the distinctiveness of activity patterns in the ventral visual cortex and parietal lobe. Critically, reduced activity was associated with enhanced representation of idiosyncratic memory traces in ventral visual cortex and precuneus. In contrast, repeated memory suppression was associated with the reduced lateral prefrontal activity, but relative intact mnemonic representations. Our results replicated most of the neural changes induced by memory retrieval and suppression immediately after learning and extended those findings to established memories after initial consolidation. Active retrieval seems to promote episode-unique mnemonic representations in the neocortex after initial encoding but also consolidation.

**Highlights:**

- Repeated retrieval strengthened consolidated memories, while repeated suppression had a modest negative effect.
- Pattern reinstatements of individual memories were detected in the visual area, parietal lobe, and hippocampus after 24 hours.
- After repeated retrieval, reduced activity amplitude was associated with increased distinctiveness of activity patterns in the ventral visual cortex and right precuneus.
- Repeated suppression was associated with the reduced lateral prefrontal activity, but unchanged mnemonic representations.

## 1. Introduction

Historically, memories were seen as more or less stable traces or engrams. After initial formation, memory traces are affected by consolidation leading to stabilization and weakening leading to forgetting (Ebbinghaus, 1885; Lashley, 1950; Müller and Pilzecker, 1900). However, contemporary research has provided ample evidence showing that memories continue to be dynamically adapted after initial encoding and thus, can be modified by external factors throughout their existence. For instance, retrieval practice can reinforce memory traces (Karpicke and Roediger, 2008), promote meaningful learning (Karpicke and Blunt, 2011), and protect memory retrieval against acute stress (Smith et al., 2016). In contrast, retrieval suppression can prevent unwanted memories to be retrieved (Anderson and Green, 2001), and reduce their emotional impact (Gagnepain et al., 2017).

Previous neuroimaging studies identified several neural changes that could explain the retrieval-mediated memory enhancement: after repeated retrieval, several studies reported decreased or increased univariate activity in frontal, parietal areas, and temporal gyrus (Eriksson et al., 2011; Gagnepain et al., 2014; Kuhl et al., 2010; Nelson et al., 2013; van den Broek et al., 2016, 2013; Wimber et al., 2011, 2008; Wing et al., 2013; Wirebring et al., 2015). More direct evidence for retrieval-induced changes in mnemonic representations came from studies that applied multivariate pattern analysis. Karlsson Wirebring and colleagues reported that less similar activity patterns in the posterior parietal region across retrieval trials are associated with subsequent better memory (Wirebring et al., 2015). Wimber and colleagues founded that targeted activity patterns are increasingly reinstated over repeated retrieval in visual areas during memory competition (Wimber et al., 2015). Most recently, Ferreira and colleagues reported retrieval-induced generalised, and episode-unique representations in parietal areas (Ferreira et al., 2019). Regarding neural changes underlying suppression-induced forgetting, compelling evidence suggested the role of prefrontal top-down regulation of the hippocampus during suppression (Anderson et al., 2004; Anderson and Hanslmayr, 2014). But only a few studies investigated neural changes in activity and/or activity patterns across repeated suppression. Depue and colleagues showed the time-specific involvement of inferior frontal gyrus and medial frontal gyrus during the suppression of emotional memory (Depue et al., 2007). Gagnepain and colleagues demonstrated the effect of suppression on visual memories may be achieved by targeted cortical inhibition of visual-related activity and activity patterns (Gagnepain et al., 2014).

Although these studies shed light upon neural changes underlying memory retrieval and suppression, all of them were based on memory modulation (i.e. retrieval and suppression) immediately after initial memory formation, except for one study that included repeated retrieval on two consecutive days (Ferreira et al., 2019). How the modulation of memory traces after initial consolidation is reflected in the neural activity and mnemonic representation, as assessed by activation patterns during subsequent retrieval is currently not well understood. Studying the neural changes underlying the modulation of initially consolidated memories can provide complementary and critical understandings of the dynamic nature of human memory. Because newly acquired memories are usually more labile compared to consolidated ones (Frankland and Bontempi, 2005) and mnemonic representations shift from the hippocampus to distributed neocortical regions following overnight sleep (Takashima et al., 2009, 2006), the effectiveness of memory modulation could be decreased and the underlying neural changes could be different. For example, a study showed that suppression of aversive memories after overnight consolidation is harder, and involved reconfigured neural pathways during suppression (Liu et al., 2016). In addition, modulation of consolidated memories may provides a clear focus on the changes of long-term memory representation, because previously reported immediate effects (i.e. changes in activity amplitude and activtiy patterns) can still be caused by short-term changes in related processes such as executive control or attention. Here, we used a two-day functional Magnetic Resonance Imaging (fMRI) design to characterize neural dynamics of initially consolidated memory. After overnight consolidation, memories were in one condition reinforced by repeated memory retrieval and in the other, weakened by repeated memory suppression. We analysed the neuroimaging data from both the modulation and the subsequent memory retrieval phase to examine neural changes at the moment when specific memory was modulated and in the final memory test in which the aftereffects of the modulation can be measured.

Based on neural findings of memory reinstatement (Chen et al., 2017; Kosslyn et al., 1997; Kuhl et al., 2010; Lee et al., 2018; O’Craven and Kanwisher, 2000; Polyn et al., 2005; Shohamy and Wagner, 2008; Wheeler et al., 2000; Wimber et al., 2015; Xue, 2018), we used both the levels of activity amplitude (i.e., univariate analysis) and activation patterns (i.e., multivariate pattern analysis) of visual area, parietal lobe, and hippocampus to characterise memory traces during memory retrieval and further examined the linear relationship between the two neural changes within the same regions. Furthermore, we adopted a novel design to disentangle perception-related neural activities associated with memory cues presented at the test and retrieval-related neural reactivation associated with reactivated mental images. One method to separate these two processes is to use two perceptual modalities (e.g. sounds as memory cues, and pictures as information to be retrieved)(Bosch et al., 2014). Here, we used highly similar visual memory cues across different memory associations. Thus, item-specific neural patterns (at least in visual areas) during retrieval more likely to be caused by retrieval-related memory reactivation instead of visual processing of memory cues.

To sum up, our primary goal is to reveal if two behavioural techniques (i.e. retrieval and suppression) can modulate initial consolidated associative memories, and if such modulation results in altered activity and/or activity patterns detected by fMRI. We first investigated the possibility that associative memories can still be modulated after 24 hours. Behaviorally, we asked whether repeated retrieval and memory suppression would oppositely strengthen or weaken original memory traces. Next, using fMRI, we examined whether retrieval and suppression would modify neural measures of memory reactivation (i.e. activity amplitude and activity pattern variability) oppositely.

## 2. Materials and Methods

### 2.1 Participants

Thirty-two right-handed, healthy young participants aged 18-35 years who were recruited from the Radboud Research Participation System finished two sessions of our experiment. They all had corrected-to-normal or normal vision and reported no history of psychiatric or neurological disease. All of them are native Dutch speakers. Two participants were excluded from further analyses due to memory performance at the chance level, three additional participants were excluded, because of excessive head motion during scanning. We used the motion outlier detection program within the FSL (i.e. FSLMotionOutliers) to detect timepoints with large motion (threshold=0.9). There are at least 20 spikes detected in these excluded participants with the largest displacement rangeing from 2.6 to 4.3 while participants included had less than 10 spikes. Neuroimaging data of one additional participant was partly used: she was excluded from the analysis of the modulation phase (Think/No-Think paradigm) due to head motion (in total 53 spike, largest displacement=5.7) only during this task, while his/her data during the other tasks were included in the analyses. Thus, data of 27 participants (16 females, age=19-30, mean=23.41, SD=3.30) were included in the analyses of the final test phase, and data of 26 participants (15 females, age=19-30, mean=23.51, SD=3.30) were included in the analyses of the modulation phase. All participants scored within normal levels when applying Dutch-versions of the Beck Depression Inventory (BDI) (Roelofs et al., 2013) and the State-Trait Anxiety Inventory (STAI) (van der Bij et al., 2003). Furthermore, because of the two-session design (24 hours’ interval), we used the Pittsburgh sleep quality index (PSQI) (Buysse et al., 1989) to assess the sleep quality between the two scanning sessions. No participants reported abnormal sleep-related behaviours or significantly reduced sleep time. The experiment was approved by, and conducted in accordance with requirements of the local ethics committee (Commissie Mensgebonden Onderzoek region Arnhem-Nijmegen, The Netherlands) and the declaration of Helsinki, including the requirement of written informed consent from each participant before the beginning of the experiment.

### 2.2 Materials

#### Locations and maps

We used 48 distinctive locations (e.g. buildings, bridges) drawn on two cartoon maps as memory cues. The maps are not corresponding to the layout of any real city in the world and participants have never been exposed to the maps before the experiment. During the task, the whole map was presented with sequentially highlighting specific locations by coloured frames as memory cues. By doing this, we kept visual processes during memory tasks largely consistent.

#### Pictures

48 pictures (24 neutral and 24 negative pictures) from the International Affective Picture System (IAPS) (Lang et al., 1997) were used in this study, and these pictures can be categorised into one of four groups: animal (e.g. cat), human (e.g. reading girl), object (e.g. clock) or scene (e.g. train station). Category information was used for the following memory-based category judgment test. All images were converted to the same size and resolution for the experiment.

#### Picture-location associations

Each picture was paired with one of the 48 map locations to form specific picture-location associations. We (W.L and J.V) carefully screened all the associations to prevent the explicit semantic relationship between picture and location (e.g. lighter at the-fire department). All 48 picture-location associations were divided into three groups for different types of modulation (See Modulation Phase). For each map, 24 locations were paired 6 pictures from each category. One-third of associations (8 associations; 2 pictures from each category) on that map were retrieval associations (i.e. “think” associations), one-third of associations were suppression associations (i.e. “no-think” associations), and remaining one-third are control associations.

### 2.3 Experiment design

#### Overview of the design

This study is a two-session fMRI experiment, with the 24 hours interval between two sessions (Figure 1A). Day1 session consists of the familiarization phase (Figure 1B), the study phase (Figure 1C), and the immediate typing test. The Day2 session consists of the second typing test, the modulation phase (Figure 1D), and the final memory test (Figure 1E). Among these phases, the familiarization, modulation, and the final memory test phase were performed in the scanner, while the study phase and two typing tests were performed in the behavioural lab.

**Figure 1.**
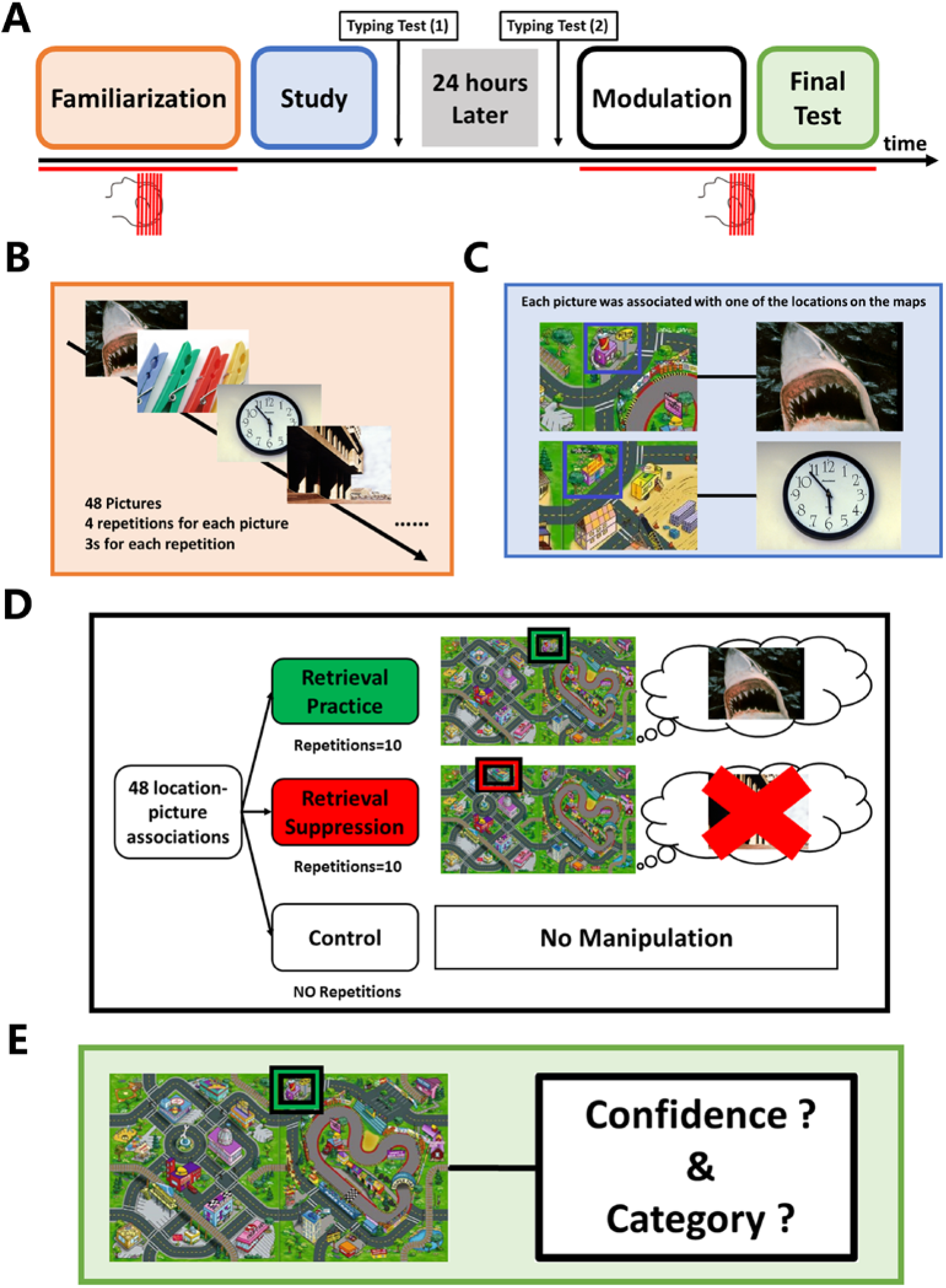
Schematic of the experiment design. **(A)** Timeline of the two-day experimental procedures. Red lines below the timeline indicate the tasks in the MRI scanner. **(B)** During the familiarization phase, all of the pictures of the to-be-remembered associations were randomly presented four times for the familiarization and estimation of picture-specific activation patterns. To keep participants focused, on each trial, they were instructed to categorise the picture shown as an animal, human, location, or object. **(C)** Study phase. Participants were trained to associate memory cues with presented pictures. **(D)** Modulation phase. After 24 hours, we used the Think/No-Think paradigm to modulate consolidated associative memories. Participants were instructed to actively retrieve associated pictures in mind (“retrieval”), or suppress the tendency to recall them (“suppression”) according to the colours of the frames (GREEN: retrieval; RED: suppression) around locations. **(E)** Final memory test phase. Participants performed the final memory test after the modulation. For each of the 48 location-picture associations, locations were presented again, and participants were instructed to report the memory confidence and categorise the picture that came to mind.

#### Familiarisation phase

To obtain the picture-specific brain responses to all 48 pictures, participants was performed initially the familiarisation phase while being scanned (Figure 1B). The second purpose of the task is to let participants become familiar with the pictures to be associated with locations later. Each picture was shown four times for 3s distributed over in total of four functional runs. The order of the presentation was pseudorandom and pre-generated by self-programmed Python code. The dependencies between the orders of different runs were minimized to prevent potential sequence-based memory encoding. To keep participants focused during the task, we instructed them to categorise the presented picture via the multiple-choice question with four options (animal, human, object, and scene). We used an exponential inter-trial intervals (ITI) model (mean=2s, minimum=1s, maximum=4s) to generate the ITIs between trials. Participants’ responses were recorded by an MRI-compatible response box.

#### Study phase

Each picture-location association was presented twice in two separate runs (Figure 1C). During each study trial, the entire map was first presented for 2.5s, then one of the 48 locations was highlighted with a BLUE frame, for 3s, and finally, the picture and its associated location were presented together for 6s. We pre-generated a pseudorandom order of the trials to minimize the similarity between the orders in familiarization and the study phase.

#### Typing test phase

Immediately after the study phase, participants performed a typing test (day1) assessing picture-location association memory. Each location was presented again (4s) in an order that differed from the study phase, and participants had maximally 60s to describe the associated picture by typing its name/description on a standard keyboard. Twenty-four hours later (day2), participants performed the typing test again in the same behavioural lab. The procedure was identical to the immediate typing test, but with a different trial order.

#### Modulation phase

The modulation phase is the first task participants performed during the Day2 MRI session. We used the think/no-think (TNT) paradigm with trial-by-trial self-report measures to modulate initially consolidated memories (Figure 1D). The same paradigm has been used in previous neuroimaging studies, and the self-report does not affect the underlying memory control process (Anderson et al., 2004; Levy and Anderson, 2012). Forty-eight picture-location associations were divided into three conditions. One-third of the associations (16 associations) were assigned to the retrieval condition (“Think”), one-third of the associations were assigned to the suppression condition (“No-Think”), and the remaining one-third of the associations were assigned to the control condition. The assignment process was counterbalanced between participants. Therefore, at the group level, for each picture-location association, the possibility of belonging to one of the three modulation conditions is around 33.3%. Associations that belong to different conditions underwent different types of modulation during this phase. Locations which belong to the control condition were not presented during this phase. For retrieval trials, locations were highlighted with a GREEN frame for 3s, and participants were instructed to recall the associated picture quickly and actively and to keep it in mind until the map disappeared from the screen. For suppression trials, locations were highlighted with a RED frame for 3s, and participants were instructed to prevent memory retrieval and to keep an empty mind. We also told the participants that they should not close their eyes or pay attention to other things outside the screen during the presentation of memory cues. After each retrieval or suppression trial, participants had up to maximum 3s to report their experience during the cue presentation. Specifically, they answered a multiple-choice question with four options (*Never, Sometimes, Often, and Always*) by pressing the button on the response box to indicate whether the associated picture entered their mind during that particular trial.

The modulation phase consisted of five functional runs (64 trials per run). In each run, 32 locations (half retrieval trials, and half suppression trials) were presented twice. Therefore, each memory cue that did not belong to the control condition was presented ten times during the entire modulation phase. Again, we pre-generated the presentation orders to prevent similar order sequences across five modulation runs. Between each trial, fixation was presented for 1-4s (mean=2s, exponential model) as ITI.

#### Final test phase

After the modulation phase, participants performed the final memory test within the scanner (Figure 1E). All 48 locations (including both the retrieval/suppression associations as well as control associations) were presented again by highlighting a specific map location with a BLUE frame. During its presentation (4s), participants were instructed to recall the associated picture covertly but as vividly as possible and keep the mental image in their mind. Critically, visual inputs during this phase were highly similar across trials because entire maps were always presented, just with different locations highlighted. Next, participants were asked to give the responses on two multiple-choice questions within 7s (3.5s for each question): (1) “how confident are you about the retrieval?” They responded with one of the four following options: Cannot recall, low confident, middle confident and high confident. (2) “Please indicate the category of the picture you were recalling?” They also had four options to choose from (Animal, Human, Object, and Scene).

### 2.4 Behavioral data analysis

#### Familiarisation phase

We did not check for the accuracy of the category judgement because there might be different opinions. However, we used individual responses to control for subjective category categorisation for the following memory performance evaluation. Specifically, if a participant consistently labelled a given picture across four repetitions as a different category compared to our predefined labels, we generated an individual-specific category label and used this category label for this picture to evaluate the responses in the final test. Otherwise, we used predefined labels to evaluate the responses.

#### Typing test

Participants’ answers were evaluated by two native Dutch experimenters (S.M and J.V) independently. The general principle is that if the answer contains enough specific information (e.g. a little black cat), to allow the experimenter to identify the picture from the 48 pictures used, it was labelled as correct. In contrast, if the answer is not specific enough (e.g. a small animal), then it was labelled as incorrect. We used Cohen’s kappa coefficient (κ) to measure inter-rater reliability. In general, κ lager than 0.81 suggests almost perfect reliability. If two accessors had different evaluations, the third accessor (W.L) determined the final result (i.e. correct or incorrect). After the immediate typing test, we only invited participants with at least 50% accuracy to the Day2 experiment. Three out of 35 recruited participants did continue on Day2 session. For the typing test 24 hours later, participants’ responses were evaluated by the same experimenters again. Based on the participants’ responses in this typing test, we identified picture-location associations that the given participant did not learn or already forgot. These associations were not considered in the following behavioural and neuroimaging analyses, because participants have no memory associations to be modulated. We calculated the average accuracies for the immediate typing test and typing test 24 hours later and investigated the delay-related decline in memory performance using a paired t-test.

#### Modulation phase

Responses during the modulation phase were analysed separately for retrieval trials and suppression trials. We first calculated the percentage of each option (never, sometimes, often, and always) chosen across 160 retrieval trials and 160 suppression trials for each participant. Next, we quantified the dynamic changes in task performance across repetitions (runs). Before the following analyses, we coded the original categorical variable using numbers (Never-1; Sometimes-2; Often-3; Always-4). For all the established picture-location associations, we calculated their average retrieval frequency rating (based on retrieval trials) and intrusion frequency rating (based on suppression trials) on each repetition. We used a repeated-measures ANOVA to model changes in retrieval and intrusion frequencies rating across repetitions to test if the repeated attempt to retrieve or suppress a memory trace would strengthen or weaken the associations respectively. Additionally, to quantify individual differences in memory suppression efficiency (Levy and Anderson, 2012), we calculated the *intrusion slope score* for each participant. Using all the intrusion rating for suppression trials, we used linear regression to calculate the slope of intrusion ratings across the ten repetitions for each participant. An increasingly negative slope score reflects better control at preventing associated memories come into awareness.

#### Final test phase

For each trial of the final test, we calculated both a subjective memory measure based on the confidence rating (1,2,3,4) and an objective memory measure based on the category judgment (correct/incorrect). Also, we recorded the reaction times (RT) for category judgments to estimate the speed of memory retrieval. To investigate the effect of types of modulation on the subjective, objective memory, and retrieval speed, we performed a repeated-measure ANOVA to detect within-participants’ differences between *RETRIEVAL ASSOCIATIONS, SUPPRESSION ASSOCIATIONS*, and *CONTROL ASSOCIATIONS*. To assess individual differences in suppression-induced forgetting, we calculated the *suppression score* by subtracting the objective memory measure of retrieval suppression associations (“no-think” items) from control association. Participants showed more forgetting as the result of suppression had more negative suppression scores.

#### Combinatory analysis of modulation and final test phase

To replicate the relationship between memory suppression efficiency during the TNT task and suppression-induced forgetting during later retrieval tests reported before (Levy and Anderson, 2012), we correlated suppression scores with intrusion slope scores across all participants. Notably, sample size (N=26) of this cross-participant correlational analysis is modest, but it is just a secondary analysis of replication.

### 2.5 fMRI data acquisition and pre-processing

#### Acquisition

MRI data were acquired using a 3.0 T Siemens PrismaFit scanner (Siemens Medical, Erlangen, Germany) and a 32 channel head coil system at the Donders Institute, Centre for Cognitive Neuroimaging in Nijmegen, the Netherlands. For each participant, MRI data were acquired in two MRI sessions (around 1 hour for each session) with 24 hours’ interval. We used three types of sequences in this study: (1) a 3D magnetization-prepared rapid gradient echo (MPRAGE) anatomical T1-weighted sequence with the following parameters: 1 mm isotropic, TE = 3.03 ms, TR = 2300 ms, flip angle = 8 deg, FOV = 256 × 256 × 256 mm; (2) Echo-planar imaging (EPI)-based multi-band sequence (acceleration factor=4) with the following parameters: 68 slices (multi-slice mode, interleaved), voxel size 2 mm isotropic, TR = 1500 ms, TE = 39 ms, flip angle =75 deg, FOV = 210 × 210 × 210 mm; (3) field map sequence (i.e. magnitude and phase images) were collected to correct for distortions (voxel size of 2 × 2 × 2 mm, TR = 1,020 ms, TE = 12 ms, flip angle = 90 deg).

During the day1 session, anatomical T1 image was acquired firstly, followed by the field map sequence. Before the four EPI-based pattern localization runs, 8 minutes of resting-state data were acquired from each participant using the same sequence parameters. Day2 session began with the field map sequence. Thereafter, we acquired six EPI-based task-fMRI runs (five runs of the modulation phase and one run of the final test phase).

#### Preprocessing of neuroimaging data

All functional runs underwent the same preprocessing steps using FEAT (FMRI Expert Analysis Tool) Version 6.00, part of FSL (FMRIB’s Software Library, www.fmrib.ox.ac.uk/fsl)(Jenkinson et al., 2012). In general, the pipeline was based on procedures suggested by Mumford and colleagues (http://mumfordbrainstats.tumblr.com) and the suggestions for Automatic Removal of Motion Artifacts (ICA-AROMA) (Pruim et al., 2015). The first four volumes of each run were removed from the 4D sequences for scanner stabilisation. The following preprocessing was applied; Motion correction using MCFLIRT (Jenkinson et al., 2002); field inhomogeneities were corrected using B0 Unwarping in FEAT; non-brain removal using BET (Smith, 2002); grand-mean intensity normalisation of the entire 4D dataset by a single multiplicative factor. We used different spatial smoothing strategies based on the type of analysis. For data used in univariate analyses, we applied a 6mm kernel. In contrast, for data used in multivariate pattern analyses, no spatial smoothing was performed to keep the voxel-wise pattern information. In addition to the default FSL motion correction algorithm, we used ICA-AROMA to further remove the motion-related spurious noise and chose the results from the “non-aggressive denoising” algorithm for the following analyses. Prior to time-series statistical analyses, highpass temporal filtering (Gaussian-weighted least-squares straight line fitting with sigma=50.0s) was applied.

Registration between all functional data, high-resolution structural data, and standard space was performed using the following steps. First, we used the Boundary Based Registration (BBR) (Greve and Fischl, 2009) to register functional data to the participant’s high-resolution structural image. Next, registration of high resolution structural to standard space was carried out using FLIRT (Jenkinson et al., 2002; Jenkinson and Smith, 2001) and was then further refined using FNIRT nonlinear registration (Andersson et al., 2007). Resulting parameters were used to align maps between native-space and standard space and back-projected region-of-interests into native space.

### 2.6 Anatomical Region-of-Interest (ROI) in fMRI analyses

Based on previous pattern reinstatement studies (Jonker et al., 2018; Lee et al., 2017, 2018; Polyn et al., 2005; Wimber et al., 2015), we hypothesised that ventral visual cortex (VVC), parietal lobe and hippocampus might carry picture-specific and category-specific information of the memory contents during retrieval. Therefore, we chose them as the ROIs in our fMRI analyses. All ROIs were first defined in the common space and back-projected into the participant’s native space for within-participant analyses using parameters obtained from FSL during registration.

We defined anatomical VVC ROI based on the Automated Anatomical Labeling (AAL) human atlas which is implemented in the WFU pickatlas software (http://fmri.wfubmc.edu/software/PickAtlas). The procedure was used before in a previous neural reactivation study conducted by Wimber and colleagues (Wimber et al., 2015). Brain regions including bilateral inferior occipital lobe, parahippocampal gyrus, fusiform gyrus, and lingual gyrus were extracted from the AAL atlas and combined to the VVC mask. The VVC mask was mainly used as the boundary to locate visual-related voxels in the following activtiy pattern analyses.

The ROIs of the hippocampus and parietal lobe (including angular gyrus (AG), supramarginal gyrus (SMG), and precuneus) were defined using a bilateral mask within the AAL provided by WFU pickatlas software. To yield better coverage of participants’ anatomies, the original mask was dilated by a factor of 2 in the software.

### 2.7 Univariate Generalized Linear Model (GLM) analyses of response amplitude

#### GLM analyses of neuroimaging data from the final test phase

To investigate how different modulations (retrieval/suppression) affect the subsequent univariate activation, we ran voxel-wise GLM analyses of the final test run. All time-series statistical analysis was carried out using FILM with local autocorrelation correction (Woolrich et al., 2001) using FEAT. In total, six regressors were included in the model. We modelled the presentation of memory cues (locations) as three kinds of regressors (duration=4s) based on their modulation history (retrieval, suppression, or control). To account for the effect of unsuccessful memory retrieval, we separately modelled the location-picture associations that participants could not recall as a separate regressor. Lastly, button press were modelled as two independent regressors (confidence and category judgment). All trials were convolved with the default hemodynamic response function (HRF) within the FSL.

We conducted two planned contrasts (retrieval vs control and suppression vs control) first at the native space and then aligned, resulting statistical maps to MNI space using the parameters from the registration. These aligned maps were used for the group-level analyses and corrected for multiple comparisons using default cluster-level correction within FEAT (voxelwise Z>3.1, cluster-level p < .05 FWER corrected). All of the contrasts were first conducted at the whole-brain level. Then, for the ROI analyses, we extracted beta values of these ROIs from the final test and compared them for the same contrasts (retrieval vs control and suppression vs control).

#### GLM analyses of neuroimaging data from the modulation phase

We ran the voxel-wise GLM analyses for each modulation run separately. In total, three regressors were included in the model. We modelled the presentation of the memory cues (location) as two kinds of regressors (duration=3s) according to their modulation instruction (retrieval or suppression). Button press was modelled as one independent regressor. In addition, if applicable, location-picture associations that our participants could not recall were modelled as a regressor. For ROI analyses, we extracted beta values of these ROIs from whole-brain maps of each modulation run separately. We investigated repetition-related changes in beta values using the Repeated ANOVA for retrieval and suppression condition separately. No multiple comparison correction was used to control for the number of ROIs involved, and we reported raw p-values for each ROI analysis.

### 2.8 Multivariate pattern analyses of brain activation patterns

#### Activity pattern estimation

All preprocessed (unsmoothed) familiarisation, modulation, and final test functional runs were modelled in separate GLMs in each participant’s native space. For each trial within familiarisation, we generated a separate regressor using the onset of picture presentation and 3s as the duration. At the same time, we generated one regressor for different button presses of the category judgment to control for the motor-related brain activity. In total, 49 regressors were included in the model. This procedure led to a separate statistical map (t-values) for each trial. Similarly, for each modulation and final test run, we generated a separate regressor using the onset of the presentation of location (memory cue) and 3s as the duration. However, button presses were not included in the model because they may potentially carry ongoing memory-related information. Also, we got a separate t map for each modulation or test trial.

#### Searchlight analysis of picture-sensitive voxels

For each participant, brain data on the familiarisation phase (i.e. pattern localisation phase) was analyzed using the searchlight method (Kriegeskorte et al., 2008, 2006) across the entire brain. More specifically, for each searchlight (centred at every voxel in the brain, a sphere with the radius of 5mm) of each participant, we trained Support Vector Classification (SVC) classifier to differentiate the activity patterns elicited by each picture (or each category) and tested its predictive power using the leave-one-run-out cross-validation. Specifically, for each trial, activity patterns within the searchlight were extracted. Since each picture was presented four times during four pattern localization runs, in total, we got four activity patterns within the searchlight for each picture. The within-participant classification was performed using the leave-one-run-out cross-validation: activity patterns of one particular run were left out as the testing dataset, and the remaining three runs were used as the training dataset to train the Support Vector Classification (SVC) model. After all the training-testing procedures, our analyses resulted in one accuracy value to represent the overall predictive power of the activity patterns within this particular searchlight. The searchlight walked through the entire brain of each participant. After the searchlight procedure, each participant yielded a classification accuracy map and each voxel within the map stored the classification accuracy of that particular searchlight sphere. To allow the group inferences of the brain regions, we performed one-sample t-tests on all of the classification accuracy maps and tested them against chance (chance level=1/48, 2%). Since we would like to identify picture-sensitive voxels within the VVC, we overlapped the voxels identified by the searchlight (p _uncorrected_<0.001) with the anatomical VVC mask. Because choosing the p _uncorrected_<0.001 as the threshold is arbitrary, we also used other thresholds (p _uncorrected_<0.05 and p _uncorrected_<0.01) to define the significant voxels and further validated our results using different threshold-dependent masks.

To further validate the localisation of picture-sensitive voxels within the VVC, we performed the ROI-based cross-participant classification based on the same pattern localisation data. Instead of performing the leave-one-run-out cross-validation, we used the leave-one-participant-out cross-validation (LOOC). Firstly, t maps for each picture and each run were transformed from native space to standard space to enable the cross-participant predictive model training and testing. Then, the identified voxels within the VVC were used as a mask to extract spatial patterns of activation. Finally, data from N-1 participants was used to train the SVC model, and the remaining participant was used to assess the model. It is notable that cross-participant classification is just the confirmatory analysis of the searchlight classification and should not be regarded as the independent analysis. The cross-participant classification was also repeated in three clusters of VVC voxels under different thresholds (p _uncorrected_<0.05, p _uncorrected_<0.01, and p _uncorrected_<0.001).

#### Pattern reinstatement analysis

The VVC voxels identified by searchlight analysis and other anatomical-defined masks (including hippocampus, AG, SMG, and precuneus) were used as the mask in the cross-task classification of memory contents. For each trial’s t-map estimated based on the final test run, we transformed it from native space to standard space. ROI-based activity patterns from both the modulation phase and final test phase were extracted using ROI masks. We performed cross-task LOOC to reveal the shared neural representation of the perception and retrieval of the same visual stimulus. Activity patterns estimated based on the pattern localisation of the N-1 training participants were used to train the SVC predictive model. We used the activity pattern during the final test evoked by the corresponding location (memory cure) of the remaining test participant, together with the trained SVC model to predict the memory content on a trial-by-trial basis. Critically, the SVC model was trained solely on the localiser data (day1), and it was applied on the final memory test (day2) without further model fitting,. Moreover, during the final test, visual input is highly similar across trials because we just highlighted each location on an identical map as the memory cue. Therefore, if the classification accuracy is higher than chance level, the classification is unlikely based on the neural responses to the memory cues only. For each ROI, we first calculated the average decoding accuracy for each participant and tested them against chance. The higher-than-chance-level decoding accuracy is the evidence for neural pattern reinstatement during memory retrieval for that particular ROI.

Without considering the modulation of each association (i.e. retrieval, suppression, or control), we demonstrated pattern reinstatement of individual memories during retrieval after 24 hours delay. Then we tested whether different modulations have different effects on the evidence (i.e. decoding accuracy or decision value(Linde-Domingo et al., 2019)) of memory reactivation. For example, if repeated retrieval increased the reactivation evidence while suppression decreased the evidence). These analyses yielded no significant results between different modulations in all ROIs investigated (*Details in Supplemental Materials; Table S1-S2*).

#### ROI-based trial-by-trial pattern variability analysis on the modulation and final test data

Representation similarity analysis (RSA) (Cohen et al., 2017) was used to calculate trial-by-trial pattern variability within particular types of test trials (e.g. recall of associations belongs to the *RETRIEVAL ASSOCIATIONS*). Given the nature of the within-participant analysis and to improve the pattern variability estimation, we based all calculations on activity patterns in the native space.

Firstly, we analysed the multivariate activation patterns of the final test. The identified VVC voxels (Figure 2A) were transformed from standard space to native space and then used as a mask to extract 3D single-trial activity patterns to 2D vectors and z-scored for the latter correlational analysis. Activation patterns of the hippcampus (Figure 2B), angular gyrus (Figure 2C), supramarginal gyrus (Figure 2D), and precuneus (Figure 2E) were extracted in the same way. After excluding all trials with incorrect memory-based category judgement, we divided the remaining trials into three conditions based on their modulation history (e.g. retrieval practice or retrieval suppression). Next, for activity patterns of trials within the same condition, we calculated neural pattern variability using Pearson correlations between all possible pairs of trials within the group (Figure 2F). The calculations led to three separate correlation matrices for three types of test trials for each participant. Finally, we used the mean value of all of the r-values located at the left-triangle to represent the neural pattern variability of the condition (higher the r-value, lower the pattern variability). All mean r-values were Fisher-r-to-z transformed before the following statistical analyses. To investigate if different modulations have different effects on memory representation during the final test, we performed two planned within-participant comparisons: [1] *RETRIEVAL ASSOCIATIONS* vs *CONTROL ASSOCIATIONS;* [2] *SUPPRESSION ASSOCIATIONS* vs *CONTROL ASSOCIATIONS*

**Figure 2.**
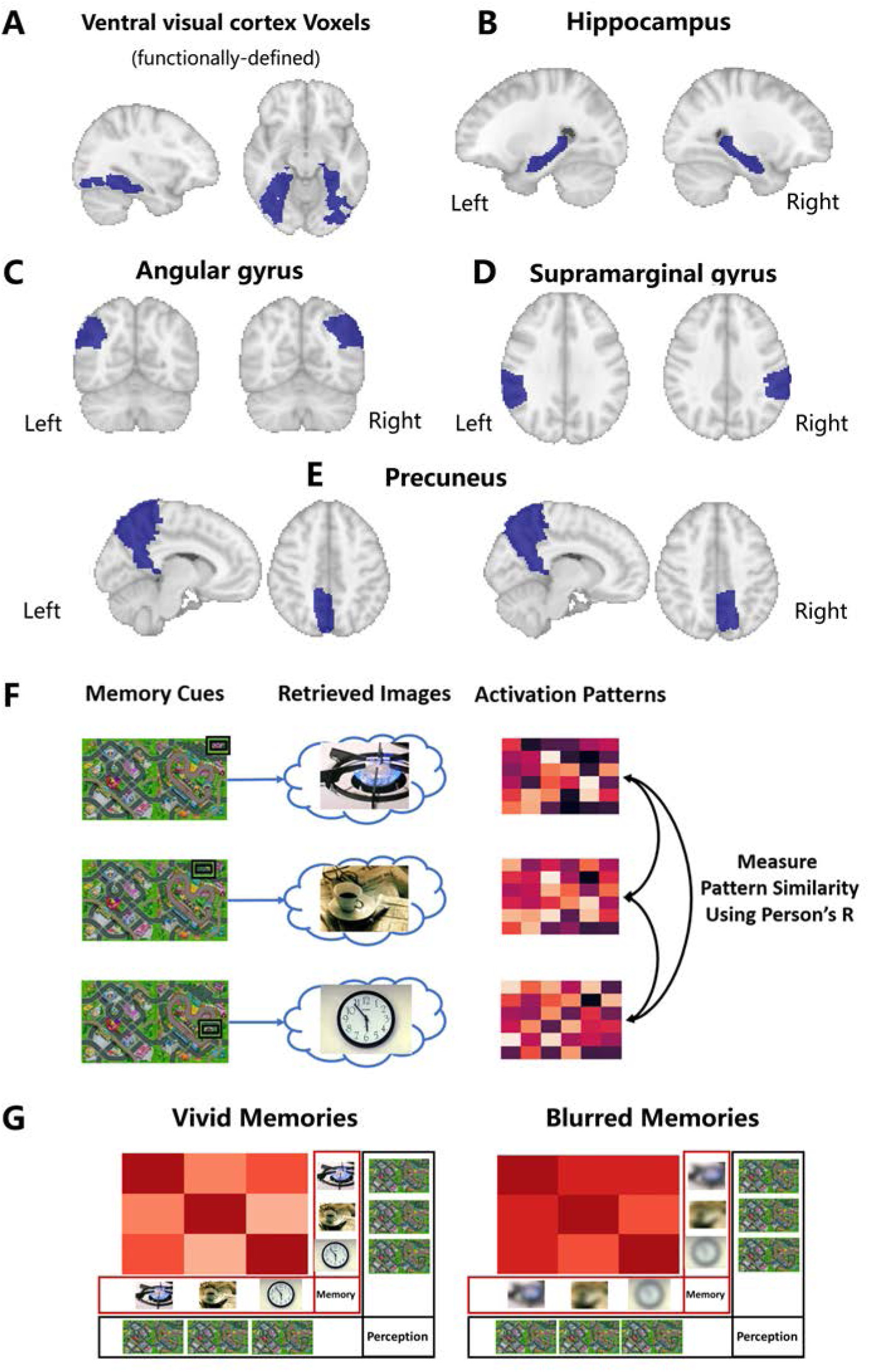
Regions-of-interest (ROI) and rationale of the activity pattern variability analysis. **(A)** Functionally-defined voxels within the ventral visual cortex (VVC). We identified voxels whose activity patterns can be used to differentiate pictures that were processed during the familiarization phase and were reactivated during successful memory retrieval during the final test. **(B)** Anatomically-defined bilateral hippocampus ROI. **(C)** Anatomically-defined bilateral angular gyrus ROI. **(D)** Anatomically-defined bilateral supramarginal gyrus ROI. **(E)** Anatomically-defined bilateral precuneus ROI. **(F)** During the final test, “mental images” were retrieved based on highly similar memory cues (different locations within maps were cued). We derived activation patterns for each memory retrieval trials based on fMRI data, and then quantify the pattern variability across trials using Person’s r. Lower the similarity measure (r-value), higher the pattern variability. **(G)** Considering the highly similar perceptional processing, vivid “mental images” during memory retrieval should be reflected in higher activity pattern variability.

Next, we used the same approach to analyse the modulation data. For each presented location, activity patterns were extracted using the same mask from five modulation runs. Similarly, within-condition (retrieval or suppression) trial-by-trial pattern variability was calculated for each condition and each run. The dynamic change was modelled using the condition by run interaction using the ANOVA analysis.

### 2.9 Data and code availability

All raw data required to reproduce all analyses and figures are uploaded onto the Donders Data Repository (https://data.donders.ru.nl/) and will be publicly available upon publication. Custom scripts used in this study will be made publicly available via the Open Science Framework (OSF) and can be requested from the corresponding authors.

## 3. RESULTS

### 3.1. Behavioural results

#### Pre-scan memory performance immediately after study and 24 hours later

During the immediate typing test (day1), 88.01% of the associated pictures were described correctly (SD= 10.87%; range from 52% to 100%). Twenty-four hours later, participants still recalled 82.15% of all associations in the second typing test (SD = 13.87%; range from 50% to 100%). Although we observed less accurate memory 24 hours later (t=4.73, p<0.001) (Figure S1), participants could still remember most location-picture associations well.

#### Behavioural performance during the modulation phase

During retrieval trials, participants reported that most associated pictures were successfully recalled (mean=84.05%, SD=11.79 %, range from 56.25% to 100%; Figure 3A). This number is close to the accuracy of the second typing test immediately before the modulation phase. Critically, we observed that with repeated attempts to retrieve, trial-by-trial retrieval frequency rating increased over repetitions, suggesting that retrieved pictures were more likely to stay stable (F [9,26]=5.77, p<0.001, η² =0.182; Figure 3B).

**Figure 3.**
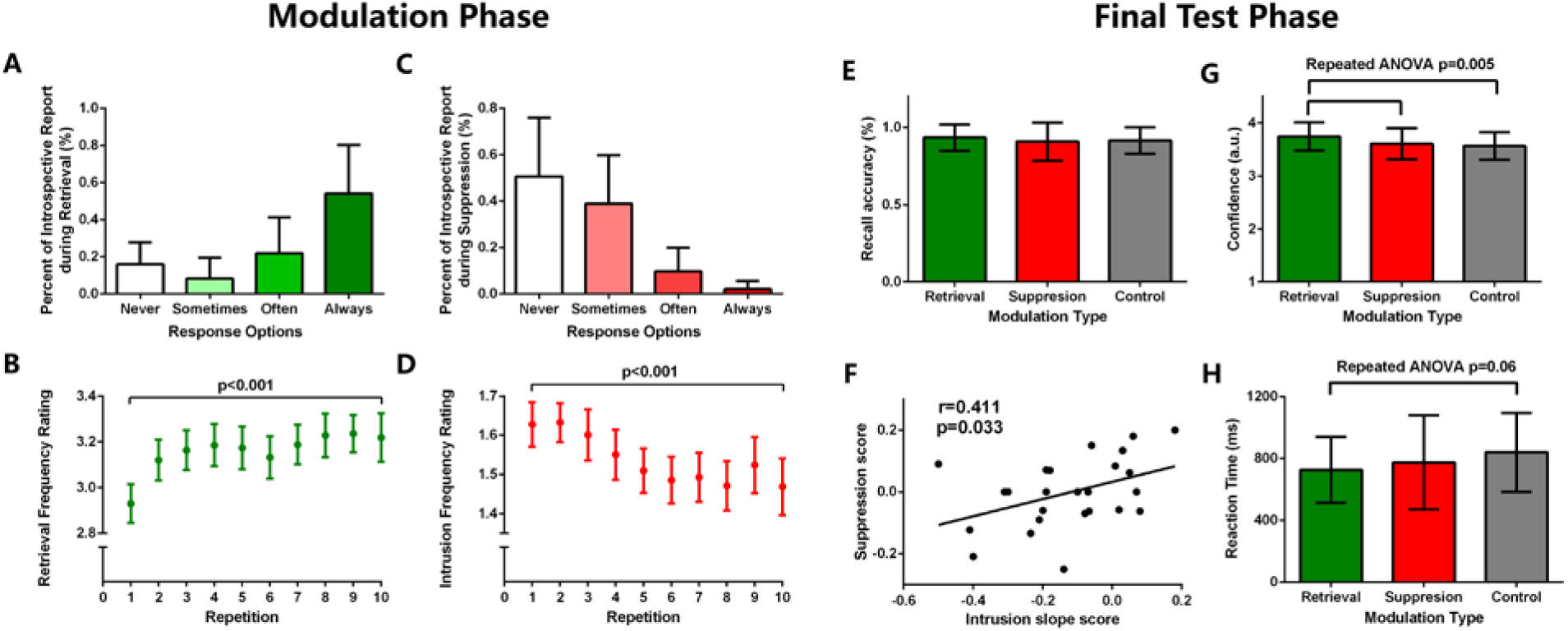
Behavioral performance during modulation and final test phase. **(A)** Percentage of the trial-by-trial introspective report during the retrieval trials. For most of the retrieval trials (mean=84.05%, SD=11.79 %), associated pictures were successfully recalled (sometimes+often+always). **(B)** With repeated retrieval attempts, associated pictures were more likely to stay in mind stably (F=5.77, p<0.001). **(C)** Percentage of the trial-by-trial introspective report during the suppression trials. During half of the suppression trials (mean=50.62%, SD=25.35%), participants successfully suppressed the tendency to recall the associated pictures (never). **(D)** As the number of repetition of suppression increase, the possibility of suppression failure declined (F=4.837, p<0.001). (E) There is no effect of retrieval or suppression on the accuracy of the categorisation during the final test (p=0.595). **(F)** Participants who are more effective in reducing suppression failures (more negative the *Intrusion Slope Score*) were the ones who show more evidence suppression-induced forgetting (more negative the *Suppression Score*). **(G)** For *RETRIEVAL ASSOCIATIONS*, participants reported higher subjective confidence compared to *SUPPRESSION ASSOCIATIONS* (t=2.172, p holm=0.07), and *CONTROL ASSOCIATIONS* (t=3.35, P _holm_=0.007). **(H)** For *RETRIEVAL ASSOCIATIONS*, participants spent less time during categorisation compared to the *CONTROL ASSOCIATIONS* (t=−2.486, P=0.02), and the effect between three conditions tend to be significant (F=2.905, p=0.06). a.u= arbitrary unit.

For the analyses of suppression trials, we excluded all location-picture associations which the participant could not describe correctly immediately before the modulation phase (i.e. Typing Test Day2). This approach controlled for individual differences in memory that could interfere with the analysis of memory suppression. On suppression trials, participants reported that they successfully suppressed the tendency to recall the associated pictures in about half of the trials (mean=50.62%, SD=25.35%, range from 4% to 92.5%; Figure 3C). As shown in think/no-think literature before (Levy and Anderson, 2012), trial-by-trial intrusion frequency rating declined from the first to the tenth repetition (F [9,26]=4.837, p<0.001, η^2^ =0.157; Figure 3D). These results suggest that participants were successful at retrieving or suppressing memory traces according to tasks instructions.

#### Memory performance during the final memory test

During the final test, participants selected, on average, the correct category (chance level=1/4) for the associated picture on 91.82% (SD = 6.05%; range from 70.83% to 100%) of the successfully recalled associations of the typing test on day2 (mean=39.43). We then examined how repeated retrieval and suppression affected memory performance. First, we compared recall accuracies between three kinds of associations (i.e. *RETRIEVAL ASSOCIATIONS, SUPPRESSION ASSOCIATIONS, and CONTROL ASSOCIATIONS*). Analysis of objective recall accuracy after modulation showed no significant main effect of *modulation* (F [2,26]=0.524, p=0.595, η^2^ =0.02; Figure 3E). Due to the lack of suppression-induced forgetting effect (lower accuracy for *SUPPRESSION ASSOCIATIONS* compared to *CONTROL ASSOCIATIONS*) at the group level, we performed a correlational analysis to associate performance during memory suppression and the final memory test. We found that participants who were more effective in suppressing intrusions (higher intrusion slope score) during the modulation phase were the ones who showed larger suppression-induced forgetting effects (r=0.411, p=0.03; Figure 3F), suggesting that successful retrieval suppression was subsequently associated with suppression-induced forgetting. This correlation was also reported before in the think/no-think literature (Levy and Anderson, 2012). Additionally, we investigated the effect of *modulation* on memory confidence and found a significant main effect (F [2,26]=5.928, p=0.005, η^2^ =0.186; Figure 3G). Post-hoc analyses revealed higher recall confidence for *RETRIEVAL ASSOCIATIONS* compared to the *CONTROL ASSOCIATIONS* (t=3.35, p _holm_=0.007) and a trend towards higher confidence compared to *SUPPRESSION ASSOCIATIONS* that just failed to reach our threshold for statistical significance (t=2.172, p _holm_=0.07). Finally, we asked if modulation affected retrieval speed indexed by the RT during the final test. Even though we did not find a significant main effect of modulation (F [2,26]=2.905, p=0.06, η^2^ =0.10; Figure 3H), recall of *RETRIEVAL ASSOCIATIONS* was faster compared to the recall of *CONTROL ASSOCIATIONS* (t=−2.486, p=0.02).

### 3.2 fMRI results

#### 3.2.1 Measuring the pattern reinstatement of individual memory during retrieval

The Support Vector Classification (SVC)-based searchlight analysis revealed brain regions including the lateral occipital cortex, fusiform gyrus, lingual gyrus, and calcarine cortex which showed picture-specific activation patterns during the perception (uncorrected p_voxel_<0.001, Figure 4A). We restricted our following activation pattern analyses to these voxels within the anatomical VVC boundary (Figure 4B). Next, we confirmed that these voxels can be used for cross-participant classification of the visual stimulus during perception. We trained the SVC based on activation patterns of N-1 participants and tested the model using the remaining subject. Results from the leave-one-out cross-validation (LOOC) confirmed these VVC voxels do enable cross-participant picture classification (mean accuracy=67.59%, SD=16.73%, one-sample t-test: t=20.37, p<0.001, Figure 4D).

**Figure 4.**
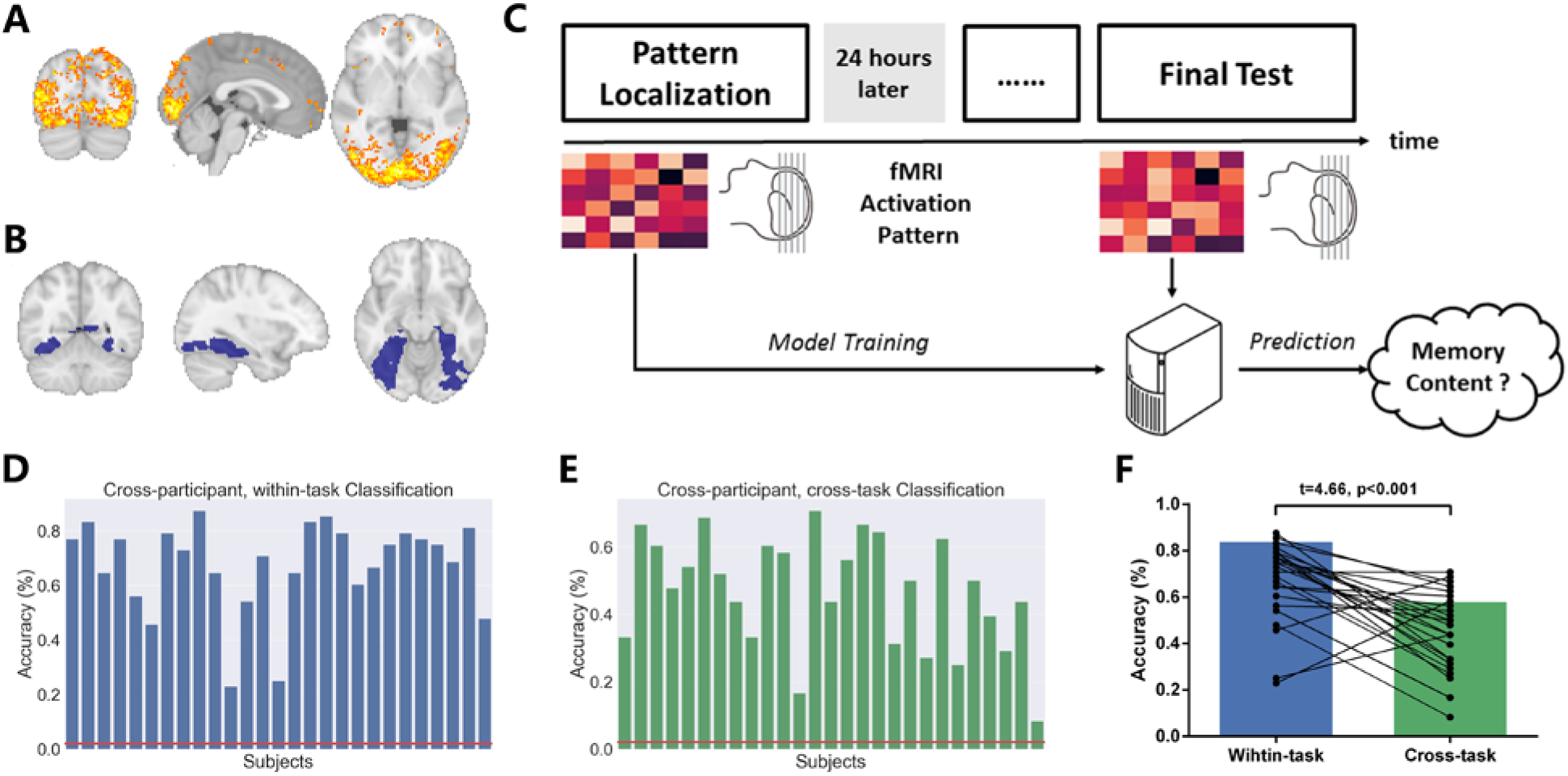
Identify picture-sensitive voxels and measure pattern reinstatement in the ventral visual cortex. **(A)** Using the searchlight method, we localised picture-sensitive voxels in brain regions included lateral occipital cortex, fusiform gyrus, lingual gyrus, calcarine cortex, postcentral and precentral gyrus, supplementary motor area, and small clusters within the medial and inferior prefrontal cortex. These voxels showed picture-specific activation patterns during the perception (uncorrected p _voxel_<0.001). **(B)** We restricted our following pattern analyses into these voxels within the ventral visual cortex (VVC) boundary by overlapping the searchlight accuracy map and anatomical-defined VVC. **(C)** fMRI activation patterns of these voxels during pattern localisation were extracted to train a classifier. Activity patterns of these voxels during the final test were further extracted and used as inputs for the classifier for different pictures. The significantly higher-than-chance level decoding of the memory contents based on activity patterns during retrieval suggests the retrieval-related pattern reinstatement in the VVC. **(D)** The classifier was first validated in a cross-participant, within-task procedure. We demonstrated that picture-sensitive voxels voxels could enable the cross-participant picture classification during perception (mean accuracy=67.59%, SD=16.73%, one-sample t-test: t=20.37, p<0.001). **(E)** The same classifier, without further model training, was used for the decoding of memory contents based on activity patterns during retrieval. Results showed that the classifier can decode the memory contents with the accuracy higher than the chance level (mean accuracy=46.84%, SD=16.82%, one-sample t-test: t=13.85, p<0.001). **(F)** We observed the significant lower classification accuracies for cross-task classification compared to the within-task classification (paired t-test: t=4.66, p<0.001).

The preceding results established that, unsurprisingly, activity patterns of voxels within the VVC carry picture-specific information during perception, we next examined if we can detect the pattern reinstatements of memory traces within the same area during the final memory test. We trained the SVC model based on the neuroimaging data from the pattern localisation phase to classify the trial-by-trial memory content in the final test (Figure 4C). Results showed that the classifiers can decode memory content based on activity patterns during the final test with an accuracy higher than chance level (mean accuracy=46.84%, SD=16.82%, one-sample t-test: t=13.85, p<0.001, Figure 4E), although the accuracy is significantly lower than the within-task classification of the perceived visual stimulus (paired t-test: t=4.66, p<0.001; Figure 4F).

We ran two control analyses to test the robustness of observed pattern reinstatement in the VVC during retrieval. We first examined the effect of arbitrary thresholds used in cluster formation on the subsequent classification of memory contents. Specifically, we used the two additional thresholds (uncorrected p _voxel_=0.01 and 0.05) to identify picture-sensitive voxels during the whole-brain searchlight analysis and confirmed that the classifications could also be performed based on picture-sensitive voxels under other thresholds (0.01 and 0.05)(Figure S2). In addition, beyond picture-specific classifications, we investigated the possibility of category-specific classifications based on brain activity patterns. All of the pictures to be associated can be categorized as one of the four following groups: animal, human, object, or location. Similarly, we localised category-sensitive voxels within the VVC (Figure S3D) and confirmed that these voxels also carry category-specific information during perception (mean accuracy=73.5%, SD=8.6%, one-sample t-test: t=29.41, p<0.001, Figure S3E). Also, activity patterns of these category-sensitive voxels during memory retrieval could enable cross-participant, cross-task classification of category during final memory test (mean accuracy=44.4%, SD=10.1%, one-sample t-test: t=10.03, p<0.001, Figure S3F).

Given the role of the hippocampus and parietal lobe in memory retrieval, we also performed the same pattern reinstatement pipeline (shown in Figure 4C) in these regions. We trained the classifier based on activity patterns of the hippocampus, angular gyrus, supramarginal gyrus, and precuneus during perception and applied it to decode memory content during retrieval. Activity patterns in these regions enabled us to perform picture-specific classification significantly higher than chance level (chance level=2%; left hippocampus: mean accuracy=7.7%, SD=3.7%, p<0.001; right hippocampus: mean accuracy=6.8%, SD=3.7%, p<0.001; left AG: mean accuracy=10.5%, SD=8.1%, p<0.001; right AG: mean accuracy=11.9%, SD=8.2%, p<0.001; left SMG: mean accuracy=9%, SD=7.4%, p<0.001; right SMG: mean accuracy=15.7%, SD=13.8%, p<0.001; left precuneus: mean accuracy=16.7%, SD=7.4%, p<0.001; right precuneus: mean accuracy=19.6%, SD=9.2%, p<0.001).

In sum, we identified picture-specific voxels within the VVC and demonstrated the pattern reinstatements of individual memory traces in these voxels during retrieval. The same pattern reinstatements were detected in anatomical-defined hippocampus, AG, SMG and precuneus. These results are the foundations of our following multivariate pattern analysis: the pattern reinstatements 24 hours after initial learning suggested that activity patterns of these regions carry mnemonic representations during retrieval.

#### 3.2.2 Repeated retrieval leads to reduced activity amplitude, but more distinct activity patterns

##### Repeated retrieval dynamically reduces the activity amplitude in the visual cortex and hippocampus

Compared to *CONTROL ASSOCIATIONS*, retrieval of *RETRIEVAL ASSOCIATIONS* was associated with less activation in medial occipital cortex, fusiform gyrus, supplementary motor area (SMA), anterior/medial cingulate cortex (MCC), left precentral gyrus, precuneus, bilateral insula, and bilateral inferior frontal gyrus (IFG) (voxelwise P_uncorrected_<0.001, p _FWE-cluster_<0.05; Figure S4A; Table S3). The VVC cluster revealed by the whole-brain analysis largely overlapped with our functional-defined VVC voxels (see Figure S4 for comparison). Our ROI analysis of these functionally-defined VVC voxels confirmed the observation: we found a reduced activity amplitude of the VVC cluster for *RETRIEVAL ASSOCIATIONS* compared to *CONTROL ASSOCIATIONS* (t=−4.8, p<0.001; Figure 5A). The whole-brain analysis did not show an effect of retrieval on the activity amplitude in hippocampal voxels under the same threshold. However, ROI-based analysis of hippocampal signal found reduced activity when retrieving *RETRIEVAL ASSOCIATIONS* compared to *CONTROL ASSOCIATIONS* (left hippocampus: t=−2.43, p=0.022; Figure 5E; right hippocampus: t=−2.18, p=0.038; Figure 5G). For six ROIs of the parietal lobe, we only found a similar retrieval-related activity reduction in the right AG (t=−2.688, df=26, p=0.012; Figure 6C) and the right precuneus (t=−2.33, df=26, p=0.027; Figure 6K), but which was not significant in the left AG (p=0.12; Figure 6A), left SMG (p=0.11; Figure 6E), right SMG (p=0.19; Figure 6G), or left precuneus (p=0.067, Figure 6I).

**Figure 5.**
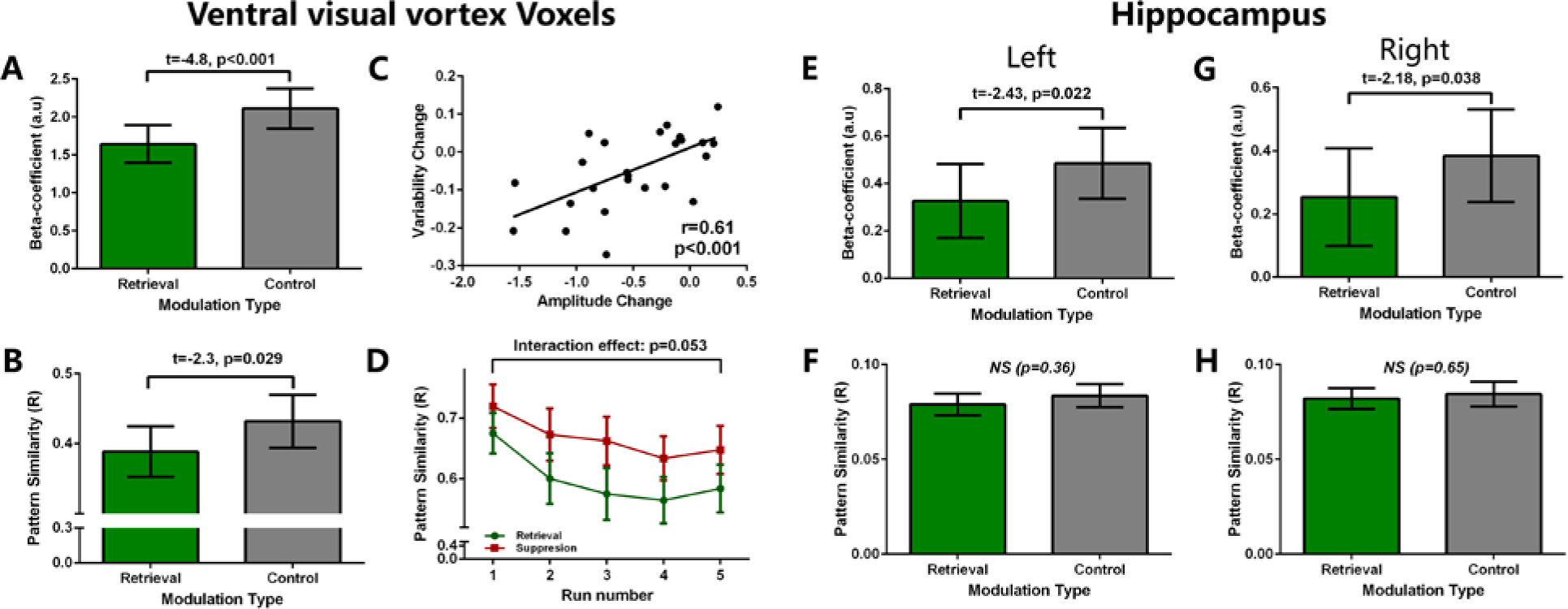
Repeated retrieval dynamically modulated activity amplitude and patterns variability. **(A)** During the final test, compared to *CONTROL ASSOCIATIONS*, *RETRIEVAL ASSOCIATIONS* was associated with lower activity amplitude in voxels within ventral visual cortex identified in the pattern reinstatement analysis. **(B)** Higher activation pattern variability (lower pattern similarity) in these VVC voxels for *RETRIEVAL ASSOCIATIONS* compared to the *CONTROL ASSOCIATIONS* during the final test (t=−2.3, p=0.029). **(C)** Across participants, the extent of activity amplitude reduction positively correlated with enhancement in pattern distinctiveness (r=0.61, p<0.001). **(D)** Dynamically enhanced pattern variability in the VVC. For both *RETRIEVAL ASSOCIATIONS* and *SUPPRESSION ASSOCIATIONS*, VVC’s pattern variability increased over repetitions during the modulation (F=11.12, p<0.001). However, repeated retrieval tends to more effectively enhance pattern variability compared to suppression (F=2.42, p=0.053). **(E)** Reduced left hippocampal activity amplitude for *RETRIEVAL ASSOCIATIONS* compared to *CONTROL ASSOCIATIONS* during the final test (t=−2.43, p=0.022). **(F)** No differences in left hippocampal activity pattern variability between *RETRIEVAL ASSOCIATIONS* and *CONTROL ASSOCIATIONS* (p=0.36) during the final test. **(G)** Reduced right hippocampal activity amplitude for *RETRIEVAL ASSOCIATIONS* compared to *CONTROL ASSOCIATIONS* during the final test (t=−2.18, p=0.038). **(H)** No differences in right hippocampal activity pattern variability between *RETRIEVAL ASSOCIATIONS* and *CONTROL ASSOCIATIONS* (p=0.65) during the final test.

**Figure 6.**
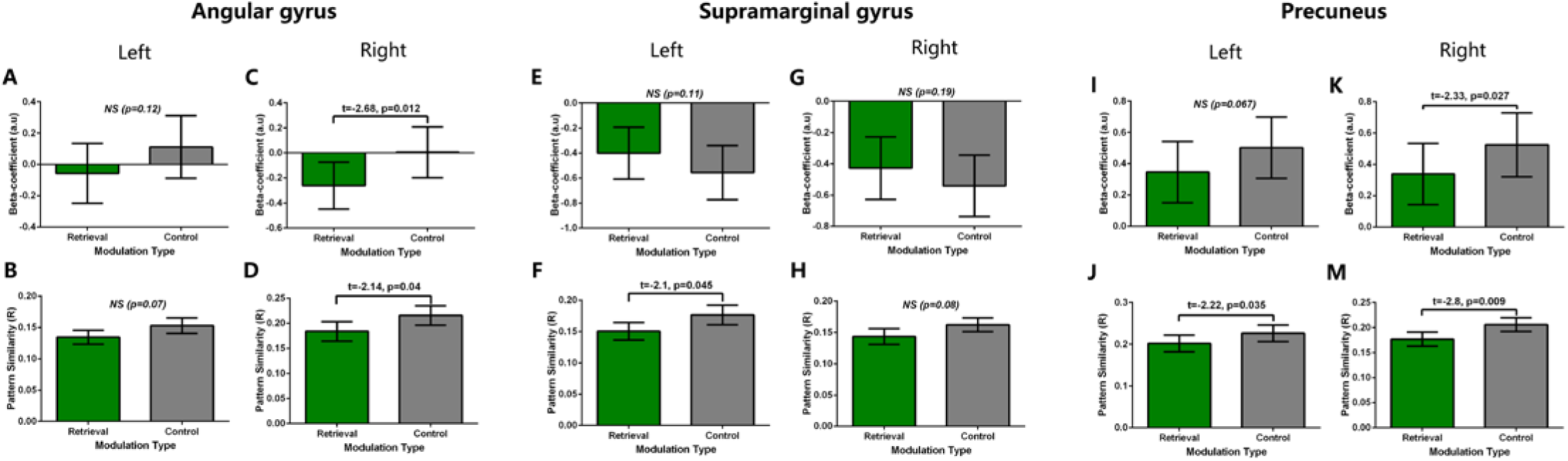
Effect of repeated retrieval on activity amplitude and patterns variability of the parietal lobe. **(A)** No differences in activity amplitude of left angular gyrus (p=0.12). **(B)** No differences in activity pattern variability of left angular gyrus (p=0.07). **(C)** Reduced activity amplitude of right angular gyrus for *RETRIEVAL ASSOCIATIONS* compared to *CONTROL ASSOCIATIONS* (t=−2.68, p=0.012). **(D)** Higher activation pattern variability (lower pattern similarity) of right angular gyrus for *RETRIEVAL ASSOCIATIONS* compared to the *CONTROL ASSOCIATIONS* (t=−2.14, p=0.04). **(E)** No differences in activity amplitude of left supramarginal gyrus (p=0.11). **(F)** Higher activation pattern variability (lower pattern similarity) of left supramarginal gyrus for *RETRIEVAL ASSOCIATIONS* compared to the *CONTROL ASSOCIATIONS* (t=−2.1, p=0.045). **(G)** No differences in activity amplitude of right supramarginal gyrus (p=0.19). **(H)** No differences in activity pattern variability of right supramarginal gyrus (p=0.08). **(I)** No differences in activity amplitude of left precuneus (p=0.067). **(J)** Higher activation pattern variability (lower pattern similarity) of left supramarginal gyrus for *RETRIEVAL ASSOCIATIONS* compared to the *CONTROL ASSOCIATIONS* (t=−2.2, p=0.035). **(K)** Reduced activity amplitude of right precuneus for *RETRIEVAL ASSOCIATIONS* compared to *CONTROL ASSOCIATIONS* (t=−2.33, p=0.027). **(M)** Higher activation pattern variability (lower pattern similarity) of left supramarginal gyrus for *RETRIEVAL ASSOCIATIONS* compared to the *CONTROL ASSOCIATIONS* (t=−2.8, p=0.009).

Next, we confirmed that the observed activity reduction is related to a linear decrease in activity with repeated retrieval using the data from the modulation phase. Specifically, we extracted the beta coefficient from these clusters for each run of the modulation phase and tested for the change in activity amplitude across runs. We found reduced VVC activity over repeated retrieval attempts (F [4, 25]=5.95, p<0.001, η^2^ =0.174). Similarly, for the bilateral hippocampus, we observed a trend toward a gradual decrease of hippocampal signal across repetitions (left hippocampus: F [4, 25]=2.39, p=0.056, η^2^ =0.087; right hippocampus: F [4, 25]=2.22, p=0.072, η^2^ =0.082). Even though we found the retrieval-related activity reduction in right AG and precuneus during the final test, we did not find the corresponding gradual decrease during modulation (right AG: F [4, 25]=0.734, p=0.571, η^2^ =0.02; right precuneus: F [4, 25]=1.88, p=0.12, η^2^ =0.05).

##### Repeated retrieval dynamically enhances the distinctiveness of activity patterns in the visual cortex, but not hippocampus

Focusing on the identified VVC voxels, parietal lobe and hippocampus, we calculated the trial-by-trial activity pattern similarity for *RETRIEVAL ASSOCIATIONS* and *CONTROL ASSOCIATIONS* separately. Results show that retrieval-related activity patterns for *RETRIEVAL ASSOCIATIONS* have increased variability in VVC compared to *CONTROL ASSOCIATIONS* (t=−2.3, df=26, p=0.029; Figure 4C). To test the robustness of increased pattern variability for *RETRIEVAL ASSOCIATIONS* in the VVC, we performed the same contrast based on (1) all associations instead of only remembered association, the VVC areas defined by (2) different thresholds and (3) category-sensitive voxels instead of picture-sensitive voxels. All control analyses yield the same result as the reported main analysis (*Figure S5-S7*). However, we did not observe a similar effect in the hippocampus (left hippocampus: t=−0.91, df=26, p=0.36, Figure 4F; right hippocampus: t=−0.456, df=26, p=0.65; Figure 4H). For six ROIs of the parietal lobe, retrieval-related enhancement of activity pattern variability was found in right AG (t=−2.148, df=26, p=0.04; Figure 6D), left SMG (t=−2.1, df=26, p=0.045; Figure 6F), left precuneus (t=−2.2, df=26, p=0.038; Figure 6J) and right precuneus (t=−2.8, df=26, p=0.009; Figure 6M). Similar trend was found in left AG (t=−1.8, df=26, p=0.07; Figure 6B) and right SMG (t=−1.79, df=26, p=0.08; Figure 6H), but failed to reach significance.

Our ROI analyses already found reduced activity amplitude, but more distinct activity patterns in VVC, right AG, and precuneus. Then we performed the correlational analysis to explore the relationship between changes in activity amplitude and changes in pattern variability across participants. We found that participants who showed a larger reduction in VVC’s activity amplitude were more likely to show a larger increase in VVC pattern variability (r=0.610, p<0.001; Figure 5C). This correlation is also significant for right precuneus (r=0.427, p=0.026), but not for right AG (r=−0.051, p=0.799).

To characterise the dynamic modulation of activity pattern variability in the VVC, we further applied the same variability analysis to each run of the modulation phase and analysed these pattern variability values using a 2×5 ANOVA (*modulation*; *run*). We saw a significant main effect of *run*, reflecting that pattern variability of the VVC increased with repetitions (F [4,25]=10.55, p<0.001, η^2^ =0.297). We also saw a main effect of *modulation*, reflecting that pattern variability of the *RETRIEVAL ASSOCIATIONS* is consistently higher than the variability of *SUPPRESSION ASSOCIATIONS* (F [1, 25]=23.77, p<0.001, η^2^ =0.487). The interaction between *modulation* and *runs* just failed to be significant (F [4, 25]=2.427, p=0.053, η^2^ =0.089; Figure 5D). This pattern of results suggests that increased pattern variability is not only the result of repetition: even though memory cues of *SUPPRESSION ASSOCIATIONS* have also been presented ten times during the modulation, repeated retrieval more effectively enhanced pattern distinctiveness compared to suppression. We applied the same dynamic modulation analysis to the right AG, left SMG, and bilateral precuneus, but found no evidence of interaction between *modulation* and *runs* (right AG: F=1.15, p=0.337; left SMG: F=0.2, p=0.938; left precuneus: F=1.81, p=0.13; right precuneus: F=0.37, p=0.82). We did not perform the same analysis to the activity patterns of the hippocampus, left AG, or right SMG because no effect was found in the final memory test.

#### 3.2.3 Retrieval suppression was associated with reduced lateral prefrontal activity

##### Weaker lateral prefrontal cortex (LPFC) activation as the result of retrieval suppression

The contrast between retrieval of *SUPPRESSION ASSOCIATIONS* and *CONTROL ASSOCIATIONS* during the final test revealed decreased activation oin one cluster in the left LPFC (x=−52,y=38, z=16, Z _peak_=4.09, size=1320 mm^3^; Figure 7A). We did not find any significant effect of retrieval suppression on hippocampal activity amplitude in the whole-brain or the ROI analysis (left hippocampus: t=−1.14, df=26, p=0.26; right hippocampus: t=−0.81, df=26, p=0.43). Also, repeated retrieval suppression was associated with reduced activity in the right AG (t=−2.07, df=26, p=0.048), but not left AG (t=−0.865, df=26, p=0.395), left SMG (t=1.214, df=26, p=0.236), right SMG (t=0.867, df=26, p=0.394), left precuneus (t=−0.77, df=26, p=0.44) or right precuneus (t=−1.13, df=26, p=0.26).

**Figure 7.**
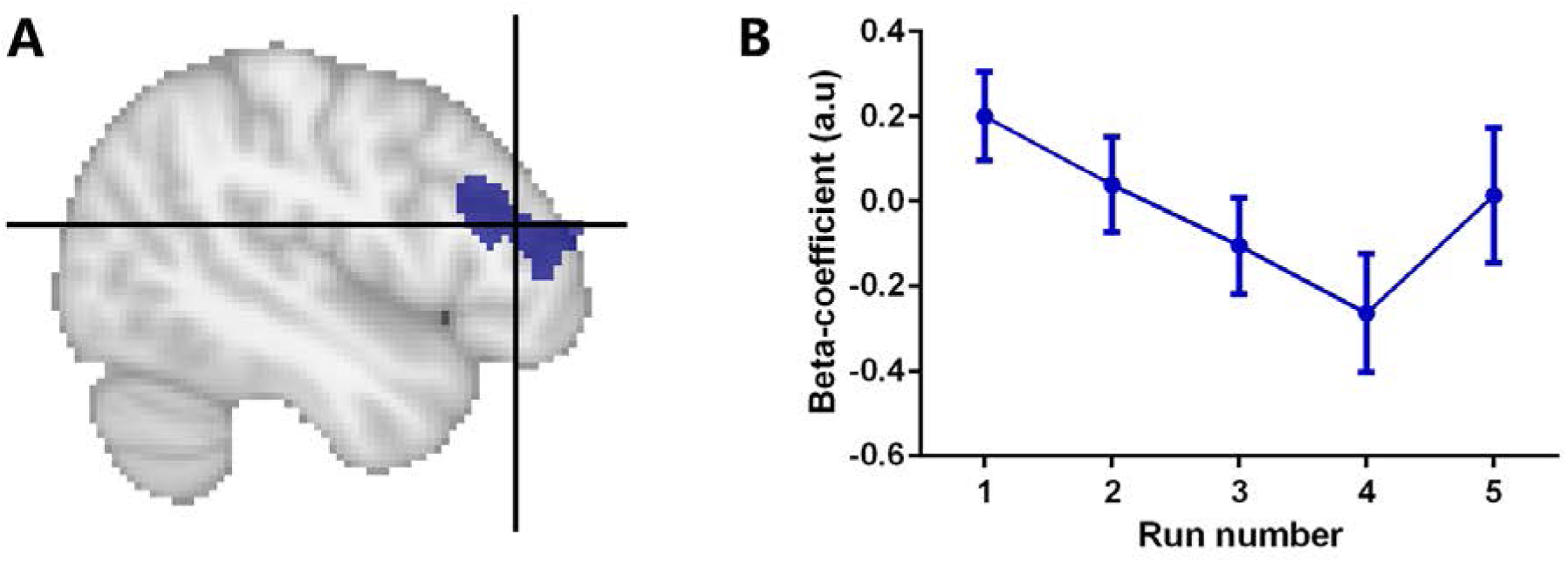
Repeated suppression disengaged lateral prefrontal cortex (LPFC) during subsequent memory retrieval. (A) During the final memory test, we found lower activity amplitude in the left LPFC for *SUPPRESSION ASSOCIATIONS* compared to *CONTROL ASSOCIATIONS*. (B) During the modulation, the activity amplitude of the same LPFC cluster tended to decreased over repetitions (from run1 to run4, p=0.03, from run1 to run5, p=0.09).

To characterise dynamical activity changes in the left LPFC, we extracted beta values from the cluster for each modulation run and found a decreased activity from the first run to the fourth run during retrieval of *SUPPRESSION ASSOCIATIONS* (F [3, 25]=2.98, p=0.036, η^2^ =0.107). However, we found an unexpected activation increase from the fourth to the fifth run, and if we combined data from all five runs, the effect failed to be significant (F [4, 25]=2.03, p=0.09, η^2^ =0.075; Figure 7B). For the right AG, we did not find any significant trend for a gradual decrease in activity during the modulation phase (F [4, 25]=1.18, p=0.323, η^2^ =0.03).

##### Intact neural representations after memory suppression

Next, we examined if retrieval suppression modulated activity patterns in the VVC, hippocampus or parietal lobe. Pattern variability analysis revealed no significant difference between *SUPPRESSION ASSOCIATIONS* and *CONTROL ASSOCIATIONS* in all regions investigated (VVC: t=−1.035, df=26, p=0.31; left hippocampus: t=−0.75, df=26, p=0.43; right hippocampus: t=−.010, df=26, p=0.92; left AG: t=0.44, df=26, p=0.663; right AG: t=0.48, df=26, p=0.63; left SMG: t=1.29, df=26, p=0.206; right SMG: t=1.15, df=26, p=0.260; left precuneus: t=−0.47, df=26, p=0.63; right precuneus: t=−1.29, df=26, p=0.2). Give the modest effect of memory suppression on final memory performance, but the strong correlation between the intrusion slope and suppression-induced forgetting, we further investigated suppression-induced changes in activity pattern variability among participants who showed strong negative intrusion slopes and (by correlation) more suppression-induced forgetting. More specifically, we used the median split method to divide the data of all participants into two groups (strong suppression group vs weak suppression group) according to their intrusion slope value and compared changes in pattern variability between groups. Our results suggested that both groups did not demonstrate differential suppression-induced changes in pattern variability for all ROIs investigated (Table S4).

## 4. DISCUSSION

Active memory retrieval is known to be a powerful memory enhancer, while memory suppression tends to prevent unwanted memories from further retrieval. Previous neuroimaging investigations of the neural effect of repeated retrieval and suppression revealed corresponding neural changes in both univariate activity analysis and multivariate activity patterns analysis. Building on these findings, we tested whether similar neural changes can be detected when modulation is delayed by 24 hours (i.e. newly acquired memories have undergone the initial consolidation). In addition, because we collected fMRI data from both the modulation phase and the final memory test, this design allowed us to perform dynamic analysis on whether the neural changes seen in the final memory test are accompanied by gradual changes during the modulation phase. Similar to previous literature (Ferreira et al., 2019), our results demonstrated that repeated retrieval of consolidated memories was associated with enhanced episode-unique mnemonic representations in the parietal lobe. Critically, our dynamic analysis provided converging evidence for the adaption of stronger mnemonic representations in visual processing areas, which were involved in the initial perception. Our results suggested that repeated retrieval of newly acquired memory and initially consolidated memory may be associated with similar neural changes.

### Repeated retrieval strengthened consolidated memories

Behaviorally, our results demonstrate that, after an initial delay of 24 hours, repeated retrieval strengthened memories further, indexed by higher recall confidence and shorter reaction times. The beneficial effect of retrieval practice on the subsequent retrieval is well established (Karpicke and Blunt, 2011; Karpicke and Roediger, 2008; Karpicke and Roediger III, 2007; Smith et al., 2016). In our study, memory accuracy was already near the ceiling level, and thus we did not find higher recall accuracy of *RETRIEVAL ASSOCIATIONS* compared to *CONTROL ASSOCIATIONS*. Corroborating the behavioural effect during the final memory test, we also found that repeated retrieval of certain memories increased their tendency to remain stable in mind during the modulation phase.

### Repeated retrieval is associated with subsequent decreasing activity amplitude

Our whole-brain univariate analysis revealed a set of brain regions including frontal, parietal (mainly precuneus) and ventral visual areas that showed decreasing activity amplitude with repeated retrieval. Activity changes in frontal and parietal areas have been reported frequently in the literature of retrieval-mediated learning/forgetting, but the direction of the reported changes are mixed. Some of the reports have found similar univariate decreases in frontal or parietal areas (Kuhl et al., 2010; Wimber et al., 2011, 2008), but others reported activity increases in these areas (Himmer et al., 2019; Nelson et al., 2013; van den Broek et al., 2016; Wirebring et al., 2015). In addition to the whole-brain analysis, our ROI analysis further showed decreased activity in the right angular gyrus. In sum, our study mainly found decreased activity in frontal and parietal areas after repeated retrieval of initially consolidated memories. Moreover, decreased activity in ventral visual areas is a novel finding. Previous studies usually used words as materials to be remembered (Nelson et al., 2013; Wimber et al., 2011, 2008; Wirebring et al., 2015), while we used pictures. One other study used also pictures and the TNT paradigm but did not reveal reliable activity changes for retrieved pictures compared to the controlled pictures (Gagnepain et al., 2014). To test the fast-consolidation hypothesis of retrieval-mediated learning (Antony et al., 2017), we further examined changes in hippocampal activity during modulation and final test. Similar to a recent report of slow hippocampal disengagement during repeated retrieval (Ferreira et al., 2019), we found dynamically decreasing hippocampal activity across repeated retrieval for initially consolidated memories. Our results, together with findings of Ferreira and colleagues, are consistent with decreasing retrieval-related hippocampal activity over the course of consolidation (Takashima et al., 2009, 2006).

### Repeated retrieval enhanced episodic-unique cortical representations

Our multivariate pattern analysis showed that compared to controls, repeated retrieval led to less similar activity patterns in ventral visual areas, and almost all parietal ROIs, including AG, SMG, and precuneus. Using a conceptually similar method, Ferreira and colleagues also reported increased item-unique activity patterns in parietal regions across two days (Ferreira et al., 2019). These results together may suggest the interaction between the effect of repeated retrieval on episodic-unique neural representations and consolidation during sleep or consolidation in general. Similar representational dissimilarity analysis has been used to analyse patterns of activity during retrieval suppression (Gagnepain et al., 2014). However, after the modulation, participants of this study only performed a visual perception task which measures repetition priming instead of a direct measure of memory. Therefore, it is impossible to directly compare the trial-by-trial pattern similarity during retrieval between *RETRIEVAL* and *CONTROL* associations.

One novel aspect of our findings is that after repeated retrieval, we found the decreased retrieval-related activity amplitude correlated with enhanced distinctiveness of activity patterns in ventral visual areas and precuneus. Our dynamic analysis of these two neural measures during modulation and subsequent memory test confirmed further that the neural changes observed during the later test are associated with dynamic adaptation of activity amplitude and pattern variability during modulation. However, this is not true for the precuneus. In general, this pattern of results is in line with our knowledge about how expectations shape brain responses. Expected stimuli reduce overall activity amplitude, a phenomenon termed “expectation suppression”(Summerfield et al., 2008; Summerfield and De Lange, 2014). At the same time, underlying activity patterns carry more visual information (de Lange et al., 2018; Kok et al., 2012). By correlating these two neural changes in the same regions, our study reported a similar phenomenon during memory retrieval. This finding suggests that the inverse relationship between overall activity amplitude and pattern-based information representation holds not only for visual expectation but also for memory retrieval. During retrieval of strengthened memories, redundant neural activity is suppressed and only the fine-grained neural patterns are reinstated, enabling more distinctive memory representations with higher fidelity.

### Retrieval suppression inhibited lateral prefrontal activity during subsequent retrieval

For *SUPPRESSION ASSOCIATIONS*, we observed lower LPFC activity amplitude, but relatively intact activity patterns in visual areas, parietal lobe, and hippocampus during subsequent retrieval. Active memory suppression during retrieval is proposed to be partially supported by inhibitory control mechanisms mediated by the lateral prefrontal cortex (Anderson and Hanslmayr, 2014; Guo et al., 2018). During retrieval suppression, LPFC is typically activated (Anderson et al., 2004; Guo et al., 2018; Levy and Anderson, 2012), but it showed gradually decreasing activity amplitudes from early suppression attempts to the later trials of suppression (Depue et al., 2007). Consistent with this pattern, we found a similar decrease in LPFC activity amplitude across suppression attempts during the modulation phase and lower activity amplitude during the subsequent retrieval. Together with the trial-by-trial intrusion frequency rating during modulation, this activity decrease across suppression attempts may suggest less inhibitory control demands when suppressing increasingly weakened memories. The observed reduction in LPFC activity during the subsequent retrieval might be a long-lasting effect of this reduced activity amplitude and suggests that modulated cognitive control allocation hampers retrieval. Another interesting observation is that we found weak evidence for suppression-induced changes in pattern reinstatement during the final memory test. Even though the involvement of the LPFC-hippocampal circuit in suppression has been examined (Anderson and Hanslmayr, 2014; Guo et al., 2018), the changes in neural representations of individual memory trace underlying suppression-induced forgetting remain less well studied. One study measured the effect of retrieval suppression on newly acquired visual memories via cortical inhibition (Gagnepain et al., 2014) and this study found that retrieval suppression reduced activity amplitude in the fusiform gyrus compared to retrieval, but the pattern was opposite to the one found in the lateral occipital complex. Effective connectivity and pattern similarity analysis suggested that top-down control mediated by the middle frontal gyrus suppressed perceptual memory traces in the visual cortex. Our study did find the comparable suppression-induced changes in activity amplitude but not mnemonic representations in the visual cortex. This may relate to the modest behavioural effects or less labile consolidated memory traces. Future studies with stronger suppression-induced forgetting effects can directly compare activity patterns between still-remembered associations and forgotten associations.

#### Limitations

Our study has two limiting aspects that should be mentioned. Firstly, given that we only found a modest effect of suppression-induced forgetting, it is difficult to interpret repeated suppression-related fMRI results. There are at least two possible reasons for this modest effect: first, due to extensive training during encoding and/or the nature of our picture-location tasks, recall accuracy for all conditions was close to ceiling level. Second, the suppression-induced forgetting effect is much smaller when memories have been consolidated (Liu et al., 2016). Thus, in line with previous studies, suppression-induced forgetting may have not emerged as the group level (Gagnepain et al., 2017; Liu et al., 2016). But we replicated two findings, confirming that our memory suppression modulation was still effective. First, when unwanted memories were suppressed repeatedly, their tendency to intrude was reduced during the TNT phase (Benoit et al., 2015; Gagnepain et al., 2017; Hellerstedt et al., 2016; Levy and Anderson, 2012; van Schie and Anderson, 2017). Second, the extent of this reduction (i.e. intrusion slope) correlated with subsequent suppression-induced forgetting effect across participants (Levy and Anderson, 2012). Given this correlation, we further compared suppression-induced neural changes between a strong and a weak suppression group, but still did not find an effect of suppression on mnemonic representations. These results may suggest that even for participants who showed suppression-induced forgetting, the underlying mnemonic representations remain intact.

A second potential limitation of our study is that we only found the effect of repeated retrieval on trial-by-trial pattern dissimilarity instead of the more direct measure of memory reactivation such as decoding accuracy or decision value (Linde-Domingo et al., 2019). It is noticeable that our pattern reinstatement analysis demonstrated that, based on activity patterns in our ROIs, the individual picture can be decoded when the classifier was trained on the localizer data (day1) before testing it on the final memory test (day2). This reinstatement laid the groundwork for our pattern dissimilarity calculation because there is evidence that these activity patterns used in the variability analysis carry indeed item-specific mnemonic information during retrieval. However, when we divided the associations into three groups (i.e. retrieval, suppression and control), we did not see the evidence that retrieval or suppression can separately modulate decoding accuracies or d values. These results may suggest that decoding accuracies or d values used here were not sensitive enough after initial consolidation, because perceptual information might already be based on the transformed representation (Xiao et al., 2017). In addition, decoding outcomes and pattern variability may associate with different aspects of mnemonic representations. Sensitive decoding depends on the reinstatement of the original representation related to the perceptual input, while pattern variability reflects episode-unique activity patterns across retrieved “mental images”. Enhanced episode-unique representations after repeated retrieval, particularly in the visual processing areas, support the following notion. Given that our memory cues (i.e. highlighted locations) are visually very similar, the changes in pattern variability in visual areas are more likely to be the result of enhanced mnemonic reinstatements instead of variability induced by visual feartures of memory cues.

## Conclusion

Taken together, our study probed the effects of repeated retrieval and suppression on initially consolidated memories. We showed that repeated retrieval dynamically reduces the activity amplitude in the visual cortex and hippocampus while enhances the distinctiveness of activity patterns in the visual cortex and parietal lobe. Moreover, reduction in activity amplitude correlated with the enhancement of episode-unique mnemonic representations in visual areas and precuneus. By contrast, repeated suppression as done here was associated with reduced lateral prefrontal activity, but intact mnemonic representations. These findings extended our understanding of neural changes underlying memory modulations from newly acquired memories to initially consolidated memories and suggest that active retrieval may strengthen episode-unique information neocortically after initial encoding and also consolidation.

## Acknowledgements

We thank Joyce van Arendonk, Joost Verchuren for the assistance of data acquisition and memory performance evaluation, Sjoerd Meijer for the memory performance evaluation, and Yingjie Shi for commenting on previous versions of the manuscript. This work was supported by a PhD fellowship from the Chinese Scholarship Council (201606990020) awarded to W.L.

## Author contributions

WL, Conception and design, Acquisition of data, Analysis and interpretation of data, Drafting or revising the article; NK, Conception and design, Analysis and interpretation of data, Drafting or revising the article; GF, Conception and design, Analysis and interpretation of data, Drafting or revising the article.

