## Supplementary Material for "Probing the neural dynamics of mnemonic representations after the initial consolidation"

**This PDF file includes:**

Supplementary Text

Figure S1-S7

Table S1-S4

Correspondence:
Wei Liu
Department of Cognitive Neuroscience
Donders Institute for Brain, Cognition and Behaviour
Radboud University Medical Centre
Trigon Building, Kapittelweg 29
6525 EN Nijmegen, The Netherlands


### 1. Robustness of neural reinstatement of individual memory in the ventral visual cortex

#### *1.1 Effect of arbitrary thresholds for cluster formation on the subsequent classifications*

During the one-sample test on all of the classification accuracy maps resulting from the searchlight analyses, we used the arbitrary threshold (uncorrected $p_{voxel}<0.001$) for the cluster formation and all of the following classification analyses were based on thresholded voxels. To account for the effect of arbitrary thresholds during the cluster formation on the following analyses, we used the two additional thresholds (uncorrected $p_{voxel}=0.01$ and 0.05) to identify picture-sensitive voxels. Repeating the cross-participant, within-task classification and cross-participant, cross-task classification, we confirmed that the classifications could also be performed based on picture-sensitive voxels under other thresholds (0.01 and 0.05)(Figure S2)

#### *1.2 Possibility of category-specific classifications.*

Beyond picture-specific classifications, we investigated the possibility of category-specific classifications based on brain activation patterns. All of the pictures to be associated can be categorized as one of the four following groups: animal, human, object, or location. Similarly, we localised category-sensitive voxels within the ventral visual cortex (VVC) (Figure S4D) and confirmed that these voxels also carry category-specific information during perception (mean accuracy=73.5%, SD=8.6%, one-sample t-test: $t=29.41$, $p<0.001$, Figure S4E). Critically, activation patterns of these *category-sensitive voxels* during memory retrieval could enable cross-participant, cross-task classification of categories of the memory (mean accuracy=44.4%, SD=10.1%, one-sample t-test: $t=10.03$, $p<0.001$, Figure S3).

**2. Comparisons of evidence for memory reactivation between *RETRIEVAL, SUPPRESSION,* and *CONTROL* associations.**

Our pattern reinstatement analysis demonstated that activtiy patterns during perception were reinstated during memory retrieval in the visual processing areas, parietal lobe, and hippocampus lob after 24 hours. Then, we investigated whether different modulation (i.e. retrieval and suppression) could modulate the memory reactivation process, indexed by different decoding outcomes for different memory associations. We used two decoding outcomes as the neural evidence for memory reactivation: [1] decoding accuracy. We analysed the predicted labels generated by the Support Vector Classification (SVC) classifier for different kinds of associations separately and calculated the average decoding accuracies of *RETRIEVAL*, *SUPPRESSION*, and *CONTROL* associations for each participant. The higher decoding accuracy for *RETRIEVAL* associations compared to *CONTROL* associations may reflect stronger direct memory reactivations induced by repeated retrieval. [2] decision value (d value)(Linde-Domingo et al., 2019). During trial-by-trial classification, together with predicted labels, we also generated the distance to the hyper-plane using the "decision_function" implemented in the SVC function. This distance measure indicates how confident the classifier was about generated predicted label at the single-trial level. We inverted each raw d value of each trial for further analysis. After the inversion, higher d value reflects more confident classification. Then we calculated the average d values for each kind of associations for each participant. Finally, we used the repeated ANOVA to compare the evidence for memory reactivation (i.e. accuracy and d value) between different associations. As shown in Table S1-S2, for all ROIs investigated, we did not found the effect of modulation on decoding outcomes.

**3. Robustness of changes in activity pattern variability for *RETRIEVAL ASSOCIATIONS*.**

We performed three control analyses to assess the robustness of retrieval-induced increase in VVC's activity pattern variability for *RETRIEVAL ASSOCIATIONS.* Firstly, we investigated if the observed change in activity pattern variability only exists for remembered associations. We reanalysed the activity pattern variability for all associations without considering the individual differences in the objective memory performance, and also found the higher activation pattern variability in the VVC for *RETRIEVAL ASSOCIATIONS* compared to *CONTROL ASSOCIATIONS* (t=3.34, p=0.002; Figure S5B). Then, we examined if the observed variability change depends on the arbitrary threshold (uncorrected $p_{voxel}<0.001$) used for picture-sensitive voxels selection. Results showed that increased activity pattern variability for *RETRIEVAL ASSOCIATIONS* could be also detected under two different thresholds ($p_{voxel}<0.01$, t=2.4, p=0.023; $p_{voxel}<0.05$, t=2.41, p=0.022; Figure S6). Finally, we further localized category-sensitive voxels within the VVC (Figure S7B) and calculated the activity pattern variability for these voxels separately for *RETRIEVAL ASSOCIATIONS* and *CONTROL ASSOCIATIONS.* The same contrast also revealed higher activation pattern variability (lower pattern similarity) for *RETRIEVAL ASSOCIATIONS* compared to the *CONTROL ASSOCIATIONS* (t=2.5, p=0.018).

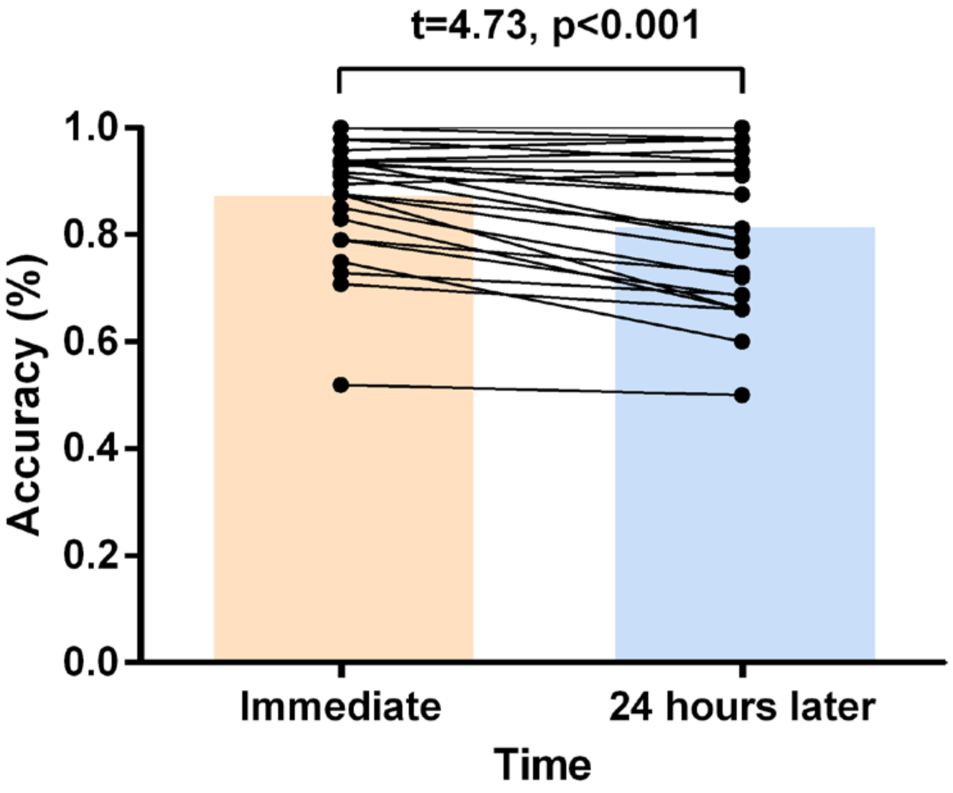


**Figure S1. Memory performance during typing test immediately after study and 24 hours later.** During the immediate typing test (day1), 88.01% of the associated pictures were described correctly (SD= 10.87%; range from 52% to 100%). Twenty-four hours later, participants could recall 82.15% of all associations (SD = 13.87%; range from 50% to 100%). Although we observed less accurate memory 24 hours later ($t=4.73$, $p<0.001$), participants could still remember most location-picture associations well.

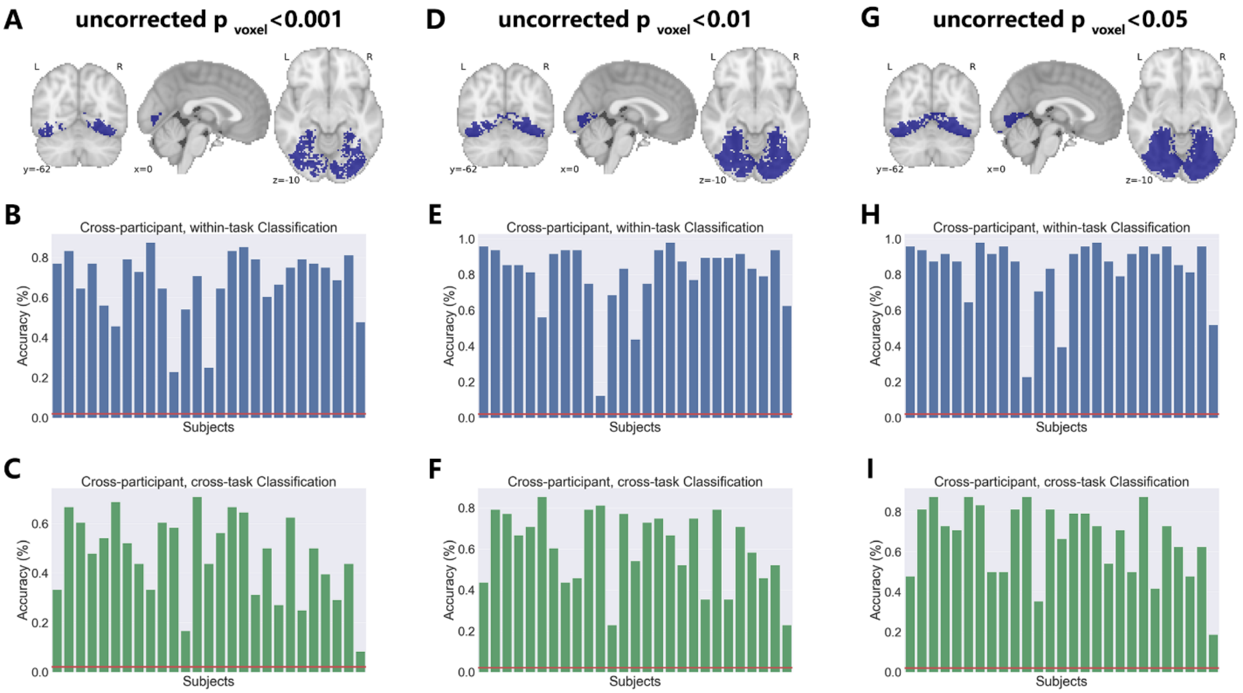


**Figure S2. Effect of different thresholds during cluster formation on the subsequent classifications. (A)** *Picture-sensitive voxels* within the ventral visual cortex identified by the searchlight method (uncorrected $p_{voxel}<0.001$). **(B)** *Picture-sensitive voxels* (uncorrected $p_{voxel}<0.001$) could enable the cross-participant picture classification during perception (mean accuracy=67.59%, SD=16.73%, one-sample t-test: t=20.37, p<0.001). **(C)** The same classifier can decode the memory contents with the accuracy higher than the chance level based on activity patterns of *Picture-sensitive voxels* (uncorrected $p_{voxel}<0.001$) during retrieval (mean accuracy=46.84%, SD=16.82%, one-sample t-test: t=13.85, p<0.001). **(D)** *Picture-sensitive voxels* within the ventral visual cortex identified by the searchlight method (uncorrected $p_{voxel}<0.01$). **(E)** *Picture-sensitive voxels* (uncorrected $p_{voxel}<0.01$) could enable the cross-participant picture classification during perception (mean accuracy=80.40%, SD=18.73%, one-sample t-test: t=21.75, p<0.001). **(F)** The same classifier can decode the memory contents with the accuracy higher than the chance level based on activity patterns of *Picture-sensitive voxels* (uncorrected $p_{voxel}<0.01$) during retrieval (mean accuracy=60.34%, SD=18.37%, one-sample t-test: t=16.50, p<0.001). **(G)** *Picture-sensitive voxels* within the ventral visual cortex identified by the searchlight method (uncorrected $p_{voxel}<0.05$). **(H)** *Picture-sensitive voxels* (uncorrected $p_{voxel}<0.05$) could enable the cross-participant picture classification during perception (mean accuracy=83.41%, SD=18.57%, one-sample t-test: t=22.78, p<0.001). **(I)** The same classifier can decode the memory contents with the accuracy higher than the chance level based on activity patterns of *Picture-sensitive voxels* (uncorrected $p_{voxel}<0.05$) during retrieval (mean accuracy=66.05%, SD=18.33%, one-sample t-test: t=18.16, p<0.001).

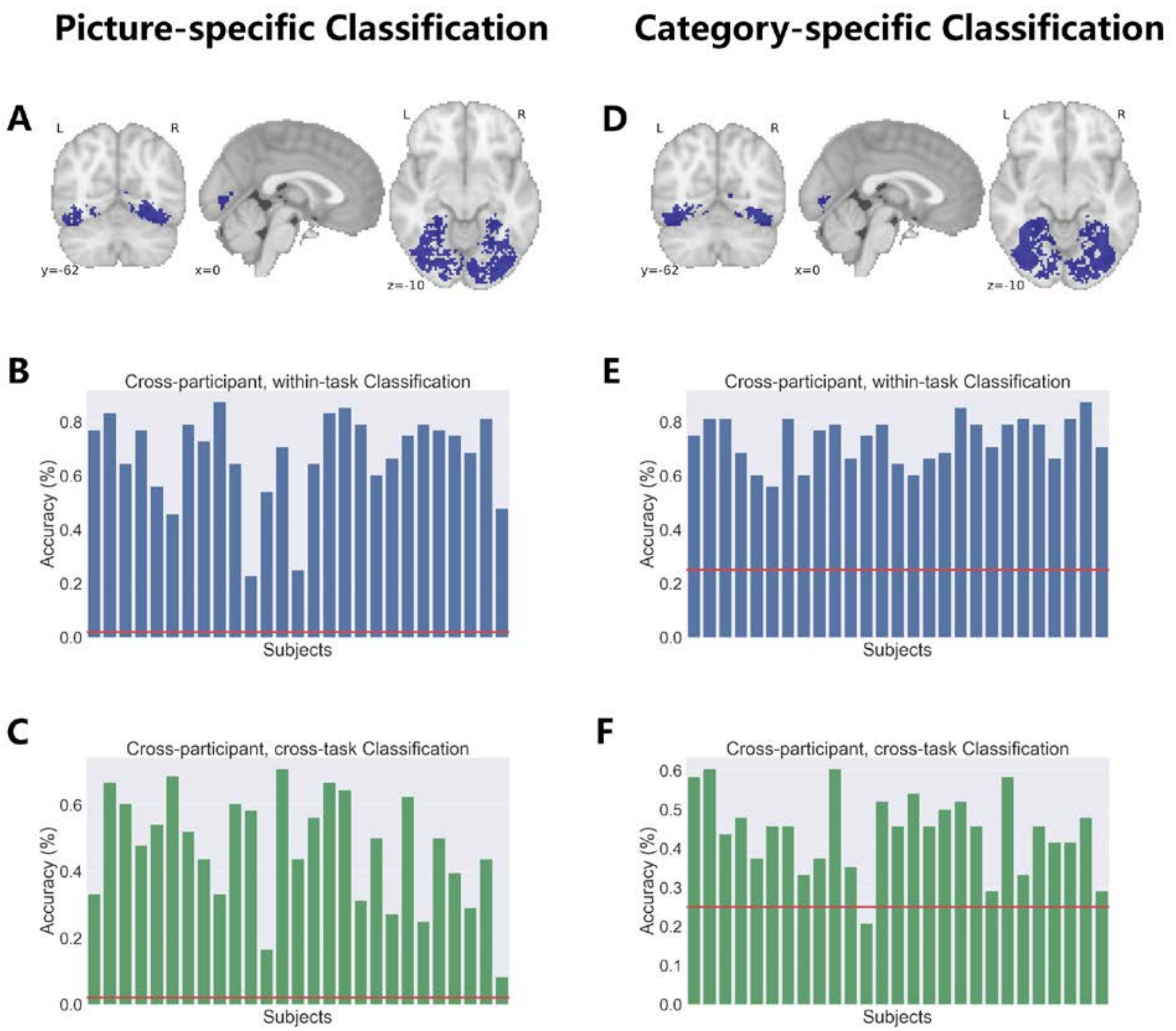


**Figure S3. Comparison between picture-specific classification and category-specific classification. (A)** *Picture-sensitive voxels* within the ventral visual cortex identified by the searchlight method. **(B)** *Picture-sensitive voxels* could enable the cross-participant picture classification during perception. **(C)** Activation patterns of *picture-sensitive voxels* during memory retrieval, together with the classifier for different pictures, could enable cross-participant, cross-task classification of memory contents. **(D)** *Category-sensitive voxels* within the ventral visual cortex identified by the searchlight method. **(E)** *Category-sensitive voxels* could enable the cross-participant picture classification during perception (mean accuracy=73.5%, SD=8.6%, one-sample t-test: t=29.41, p<0.001). **(F)** Activation patterns of *category-sensitive voxels* during memory retrieval, together with the classifier for different categories, could enable cross-participant, cross-task classification of memory contents (mean accuracy=44.4%, SD=10.1%, one-sample t-test: t=10.03, p<0.001).

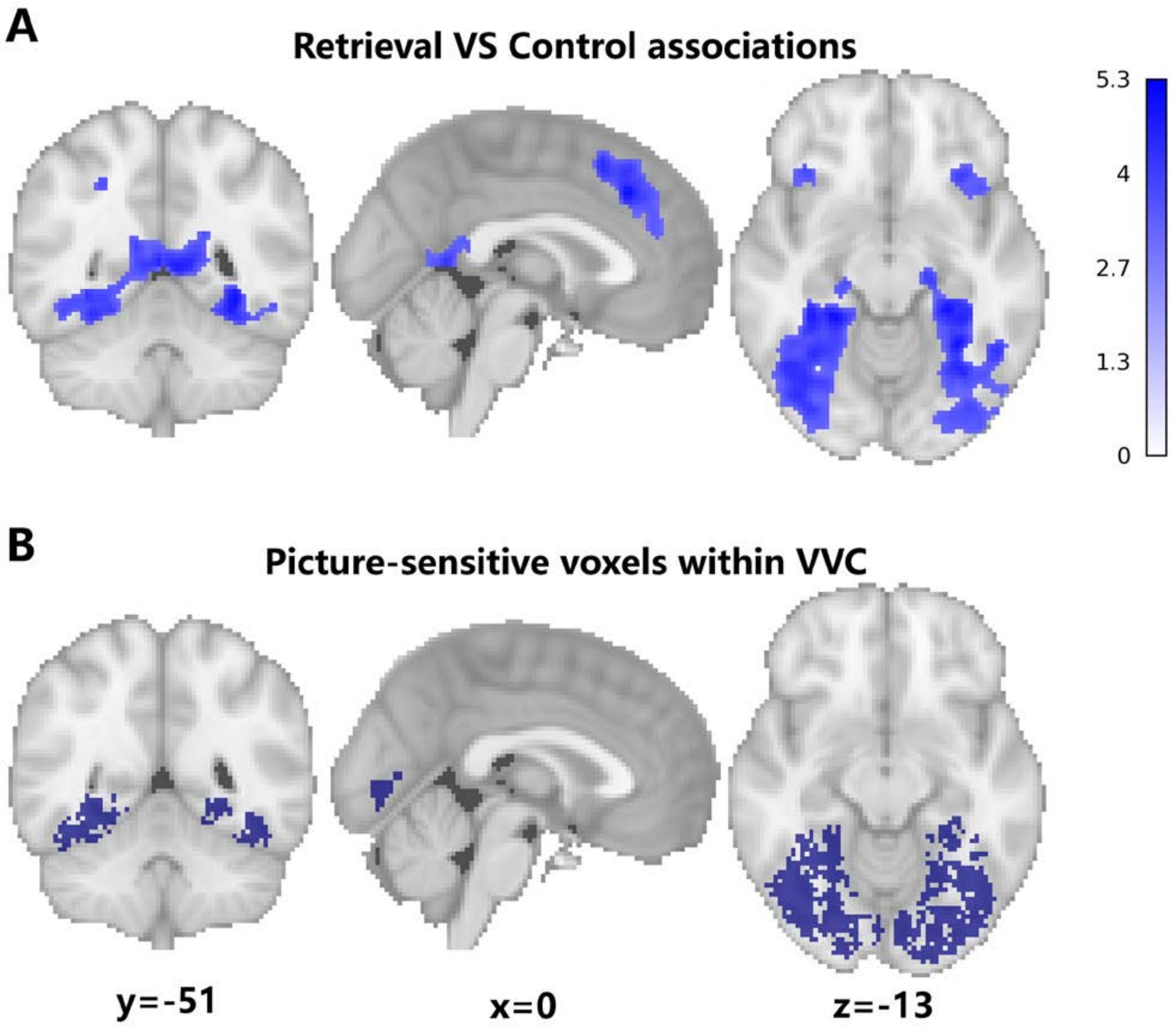


**Figure S4. Similar spatial pattern between areas showed reduced activity amplitude and picture-sensitive voxels within the ventral visual cortex. (A)** Brain regions showed less activation when retrieved *RETRIEVAL ASSOCIATIONS* compared to *CONTROL ASSOCIATIONS.* Compared to *CONTROL ASSOCIATIONS*, retrieval of *RETRIEVAL ASSOCIATIONS* was associated with less activation in medial occipital cortex, fusiform gyrus, supplementary motor area (SMA), anterior/medial cingulate cortex (MCC), precuneus, bilateral insula, and bilateral inferior frontal gyrus (IFG) (voxelwise $_{\text{uncorrected p}}<0.001$, p $_{\text{FWE-cluster}}<0.05$). **(B)** Searchlight analysis identified picture-sensitive voxels within the ventral visual cortex.

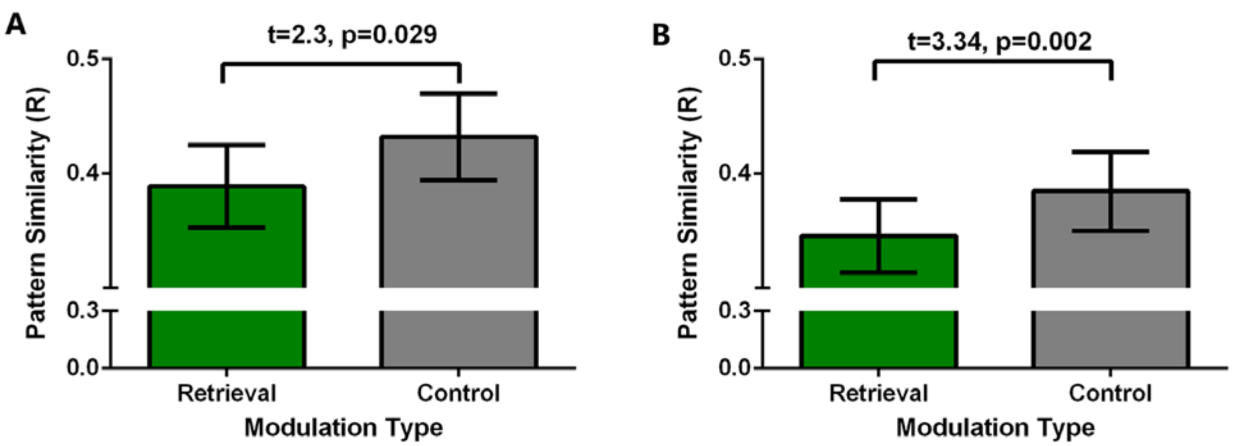


**Figure S5. Activity pattern variability analyses using all associations or only remembered associations. (A)** Using only the activation patterns of remembered associations, we found higher activation pattern variability (lower pattern similarity) in the VVC for *RETRIEVAL ASSOCIATIONS* compared to the *CONTROL ASSOCIATIONS* (t=2.3, p=0.029). **(B)** Using activation patterns of all associations, we also found the higher activation pattern variability in the VVC for *RETRIEVAL ASSOCIATIONS* (t=3.34, p=0.002).

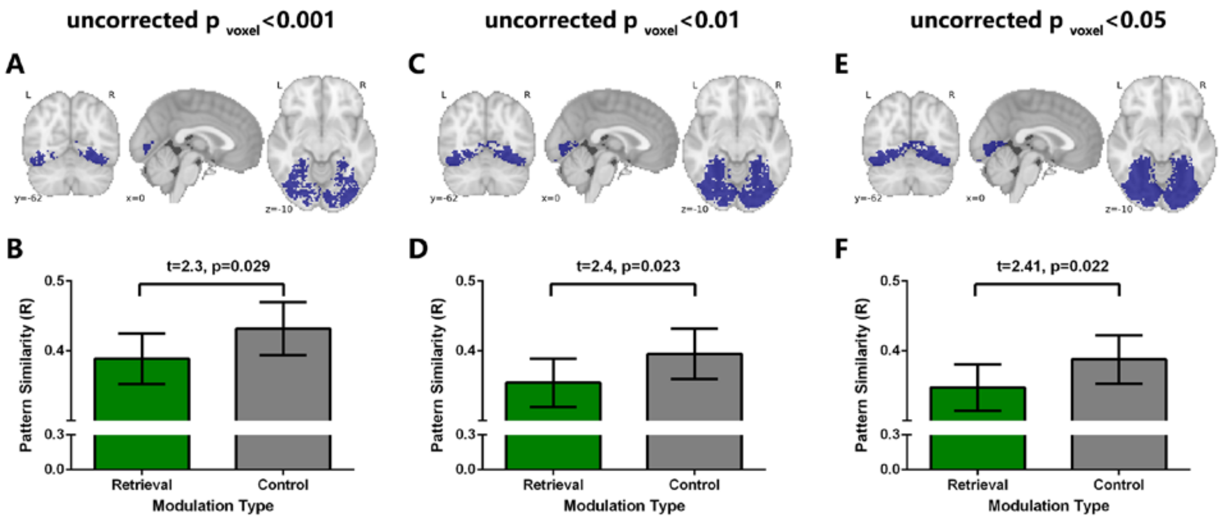


**Figure S6. Activity pattern variability analyses based on different cluster-formation thresholds. (A)** *Picture-sensitive voxels* within the ventral visual cortex identified by the searchlight method (uncorrected $p_{voxel}<0.001$). **(B)** Based on the VVC region-of-interest formed by threshold $p<0.001$, we found higher activation pattern variability (lower pattern similarity) in the VVC for *RETRIEVAL ASSOCIATIONS* compared to the *CONTROL ASSOCIATIONS* (t=2.3, p=0.029). **(C)** *Picture-sensitive voxels* within the ventral visual cortex identified by the searchlight method (uncorrected $p_{voxel}<0.01$). **(D)** Based on the VVC region-of-interest formed by threshold $p<0.01$, we also found higher activation pattern variability (lower pattern similarity) in the VVC for *RETRIEVAL ASSOCIATIONS* compared to the *CONTROL ASSOCIATIONS* (t=2.4, p=0.023). **(E)** *Picture-sensitive voxels* within the ventral visual cortex identified by the searchlight method (uncorrected $p_{voxel}<0.05$). **(F)** Based on the VVC region-of-interest formed by threshold $p<0.01$, we also found higher activation pattern variability (lower pattern similarity) in the VVC for *RETRIEVAL ASSOCIATIONS* compared to the *CONTROL ASSOCIATIONS* (t=2.41, p=0.022).

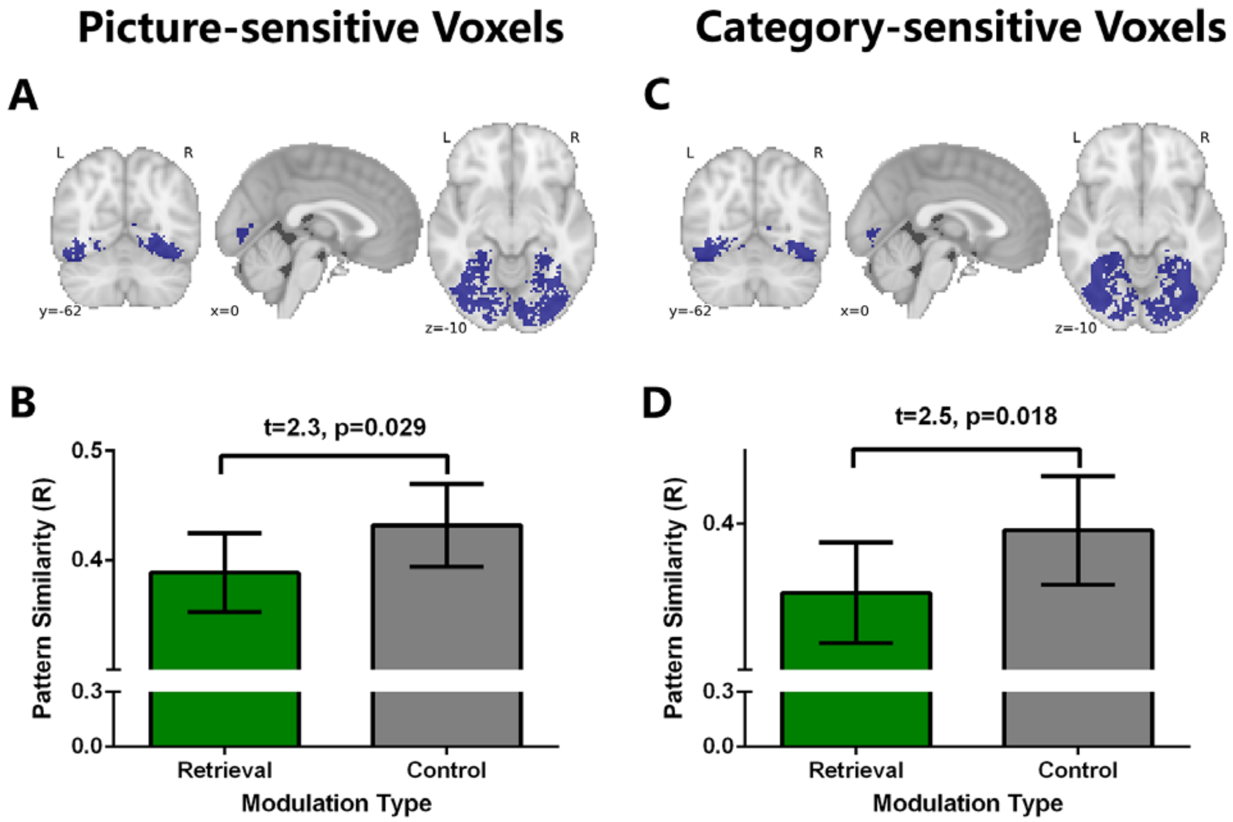


**Figure S7. Activity pattern variability analyses based on picture-sensitive voxels or category-sensitive voxels.** **(A)** *Picture-sensitive voxels* within the ventral visual cortex identified by the searchlight method. **(B)** Based on activation patterns of *Picture-sensitive voxels* within the ventral visual cortex, we found higher activation pattern variability (lower pattern similarity) in the VVC  for *RETRIEVAL ASSOCIATIONS* compared to the *CONTROL ASSOCIATIONS* (t=2.3, p=0.029). **(C)** *Category-sensitive voxels* within the ventral visual cortex identified by the searchlight method. **(D)** Based on activation patterns of *Category-sensitive voxels* within the ventral visual cortex, we also found higher activation pattern variability (lower pattern similarity) in the VVC  for *RETRIEVAL ASSOCIATIONS* compared to the *CONTROL ASSOCIATIONS* (t=2.5, p=0.018).

**Table S1 Comparisons of decoding accuracies between different modulations**

| Region of Interest (ROI) | Retrieval association (%) (mean(SD)) | Suppression association (%) (mean(SD)) | Control association (%) (mean(SD)) | F | p | $\eta^2$ |
|---|---|---|---|---|---|---|
| Ventral Visual Cortex | 44.2(17.8) | 47.5(18.4) | 48.8(19.5) | 1.64 | 0.20 | 0.01 |
| Left hippocampus | 7.9(6.2) | 8.1(7.8) | 7.2(6.4) | 0.13 | 0.87 | 0.004 |
| Right hippocampus | 7.9(7.9) | 7.2(7.7) | 5.6(5) | 0.73 | 0.48 | 0.02 |
| Left AG | 10(8.4) | 11.1(10) | 10.4(12) | 0.15 | 0.85 | 0.002 |
| Right AG | 12(9.9) | 12.7(9.9) | 10.9(11.8) | 0.35 | 0.70 | 0.005 |
| Left SMG | 9.5(9.9) | 8.3(7.6) | 9.3(10.6) | 0.19 | 0.82 | 0.003 |
| Right SMG | 15.5(15.6) | 14.4(14.2) | 17.4(16.7) | 0.79 | 0.45 | 0.007 |
| Left precuneus | 18.3(11.1) | 14.6(10.1) | 17.4(12.3) | 0.93 | 0.39 | 0.02 |
| Right precuneus | 20.4(11) | 19.4(11.9) | 19(12.7) | 0.15 | 0.85 | 0.002 |

SMG=supramarginal gyrus; AG=angular gyrus.

**Table S2 Comparisons of decoding decision value (d) between different modulations**

| **Region of Interest (ROI)** | **Retrieval association (d)** (mean(SD)) | **Suppression association (d)** (mean(SD)) | **Control association (d)** (mean(SD)) | **F** | **p** | **η²** |
|---|---|---|---|---|---|---|
| Ventral Visual Cortex | 39.27(5.06) | 40.80(5.17) | 39.75(6.42) | 1.62 | 0.20 | 0.01 |
| Left hippocampus | 28.08(3.85) | 27.62(3.52) | 27.66(4.24) | 0.12 | 0.88 | 0.002 |
| Right hippocampus | 27.89(4.16) | 27.11(3.73) | 26.71(4.01) | 0.66 | 0.51 | 0.01 |
| Left AG | 26.36(7.47) | 27.06(8.01) | 26.02(7.57) | 0.26 | 0.76 | 0.003 |
| Right AG | 27.03(8.07) | 27.21(6.21) | 26.94(8.21) | 0.02 | 0.97 | 0 |
| Left SMG | 27.70(6.74) | 28.05(7.00) | 27.63(6.87) | 0.12 | 0.88 | 0.001 |
| Right SMG | 32.67(7.60) | 31.73(6.29) | 32.61(8.62) | 0.47 | 0.62 | 0.003 |
| Left precuneus | 30.72(6.11) | 31.37(6.83) | 31.34(4.15) | 0.21 | 0.81 | 0.003 |
| Right precuneus | 30.4(5.05) | 30.43(7.04) | 31.89(4.28) | 1.30 | 0.28 | 0.016 |

SMG=supramarginal gyrus; AG=angular gyrus.

**Table S3. Brain regions showed less activation when retrieved *RETRIEVAL ASSOCIATIONS* compared to *CONTROL ASSOCIATIONS.***

| Brain region | Hemisphere | Peak MNI coordinates | Cluster size ($mm^3$) | Cluster mean |
|---|---|---|---|---|
| IFG | L | -36 42 8 | 184 | -3.23 |
| Left DLPFC | L | -46 46 6 | 224 | -3.20 |
| Precuneus | L | -8 -80 46 | 360 | -3.23 |
| Precentral gyrus | L | -38 2 38 | 3024 | -3.56 |
| Insula | R | 32 26 -6 | 9528 | -3.61 |
| Insula | L | -30 26 -2 | 10936 | -3.69 |
| ACC/MCC/SMA | R/L | 2 28 48 | 11712 | -3.68 |
| Medial occipital cortex/Fusiform gyrus | R/L | 32 -46 -8 | 71664 | -3.66 |

IFG= Inferior Frontal Gyrus; DLPFC= Dorsolateral Prefrontal Cortex; ACC= Anterior Cingulate Cortex; MCC= Middle Cingulate Cortex; SMA= Supplementary Motor Area

**Table S4 Comparison of pattern variability change between participants who showed strong and weak suppression**

| Region of Interest (ROI) | Pattern Variability change (Strong) (mean(SD)) | Pattern Variability change (Weak) (mean(SD)) | t | p | d |
|---|---|---|---|---|---|
| Ventral Visual Cortex | -0.02(0.06) | -0.006(0.08) | -0.644 | 0.526 | -0.253 |
| Left hippocampus | -0.005(0.02) | 0.009(0.01) | -0.612 | 0.546 | -0.246 |
| Right hippocampus | -0.0005(0.03) | -0.0004(0.02) | -0.01 | 0.99 | -0.005 |
| Left AG | -0.015(0.04) | 0.013(0.06) | -1.288 | 0.21 | -0.505 |
| Right AG | -0.03(0.09) | 0.02(0.08) | -1.57 | 0.129 | -0.616 |
| Left SMG | -0.007(0.04) | -0.015(0.05) | 0.4 | 0.693 | 0.157 |
| Right SMG | -0.012(0.06) | -0.011(0.04) | -0.03 | 0.973 | -0.013 |
| Left precuneus | -0.02(0.04) | 0.01(0.05) | -1.82 | 0.073 | -0.734 |
| Right precuneus | -0.028(0.06) | 0.002(0.04) | -1.35 | 0.189 | -0.53 |

Pattern Variability change=pattern variability of suppression associations minus variability of control associations; Strong=the group of participants who showed stronger suppression (i.e. more negative suppression slope); weak= the group of participants who showed weaker suppression (i.e. less negative suppression slope); SMG=supramarginal gyrus; AG=angular gyrus.
